## Supplementary Information for "Microinterfaces in bicontinuous hydrogels guide rapid 3D cell migration"

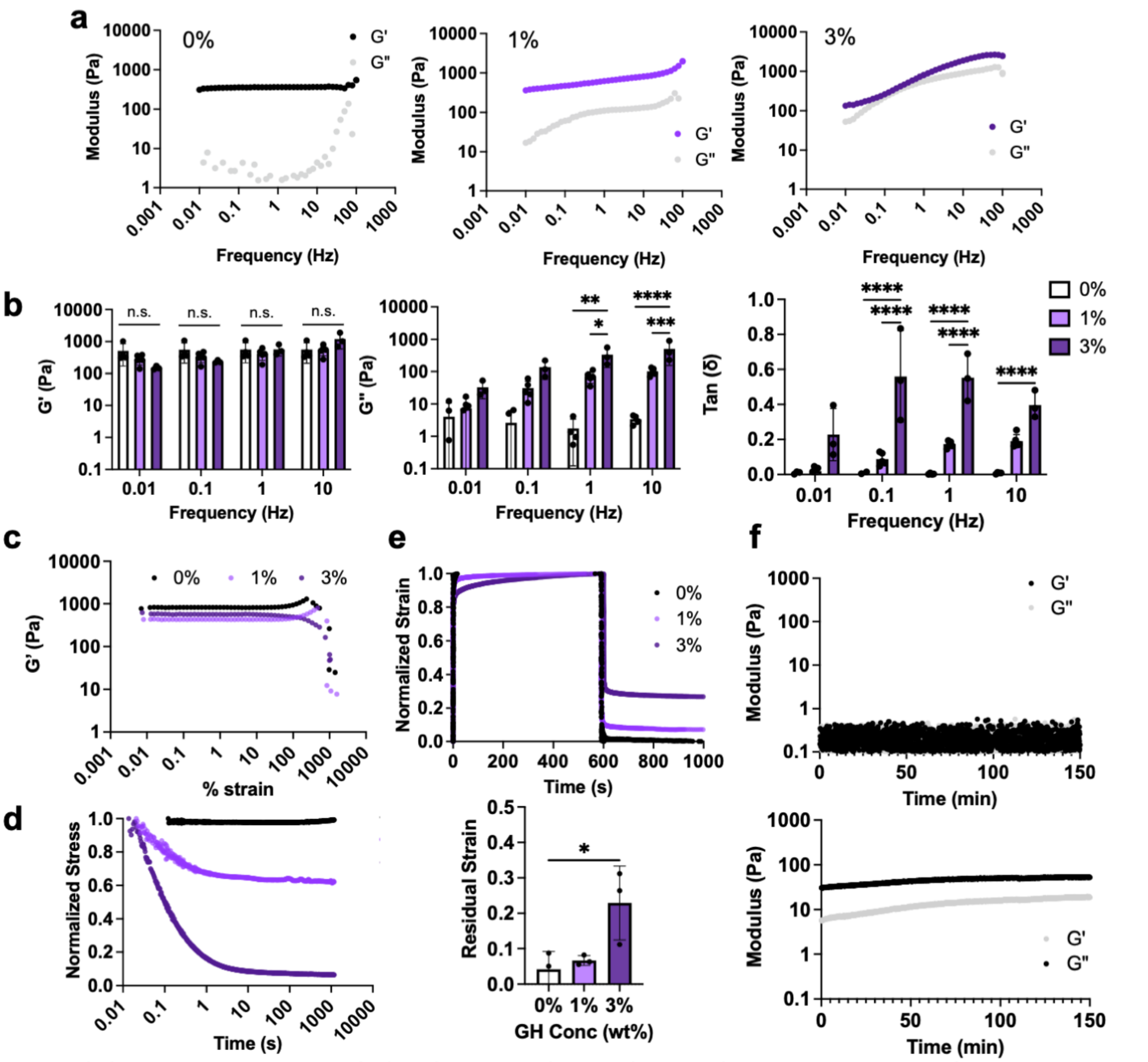


**Extended Data Fig. 1: Bicontinuous hydrogels are viscoplastic and stress-relaxing. a**, Representative frequency sweeps (0.01–100 Hz, 1% strain). **b,** Rheological measurements extrapolated from frequency sweeps across different frequencies (0.01, 0.1, 1, 10 Hz, 1% strain) of storage modulus (left panel), loss modulus (middle panel), and tan (delta) (right panel). n= 3-5 hydrogels per condition, n.s. indicates no statistical significance, *$p \leq .05$, **$p \leq .01$, ***$p \leq .001$, ****$p \leq .0001$, two-way ANOVA with Tukey post hoc. **c,** Representative strain sweeps (1 Hz, 0.001 to 1000% strain). **d,e,** Representative stress relaxation (**d**, 10% strain) and creep-recovery (**e**-top panel, 100 Pa) studies and quantification of residual strain from creep-recovery studies (**e**-bottom panel). n= 3 hydrogels per condition. *$p \leq .05$, one-way ANOVA with Tukey post hoc. **f,** Representative gelation kinetics (1 Hz, 1% strain) of 5 wt% gelatin without enzymatic crosslinker or GH (top panel) and of 5wt% gelatin with GH but without enzymatic crosslinker (bottom panel). Data are mean ± s.d.

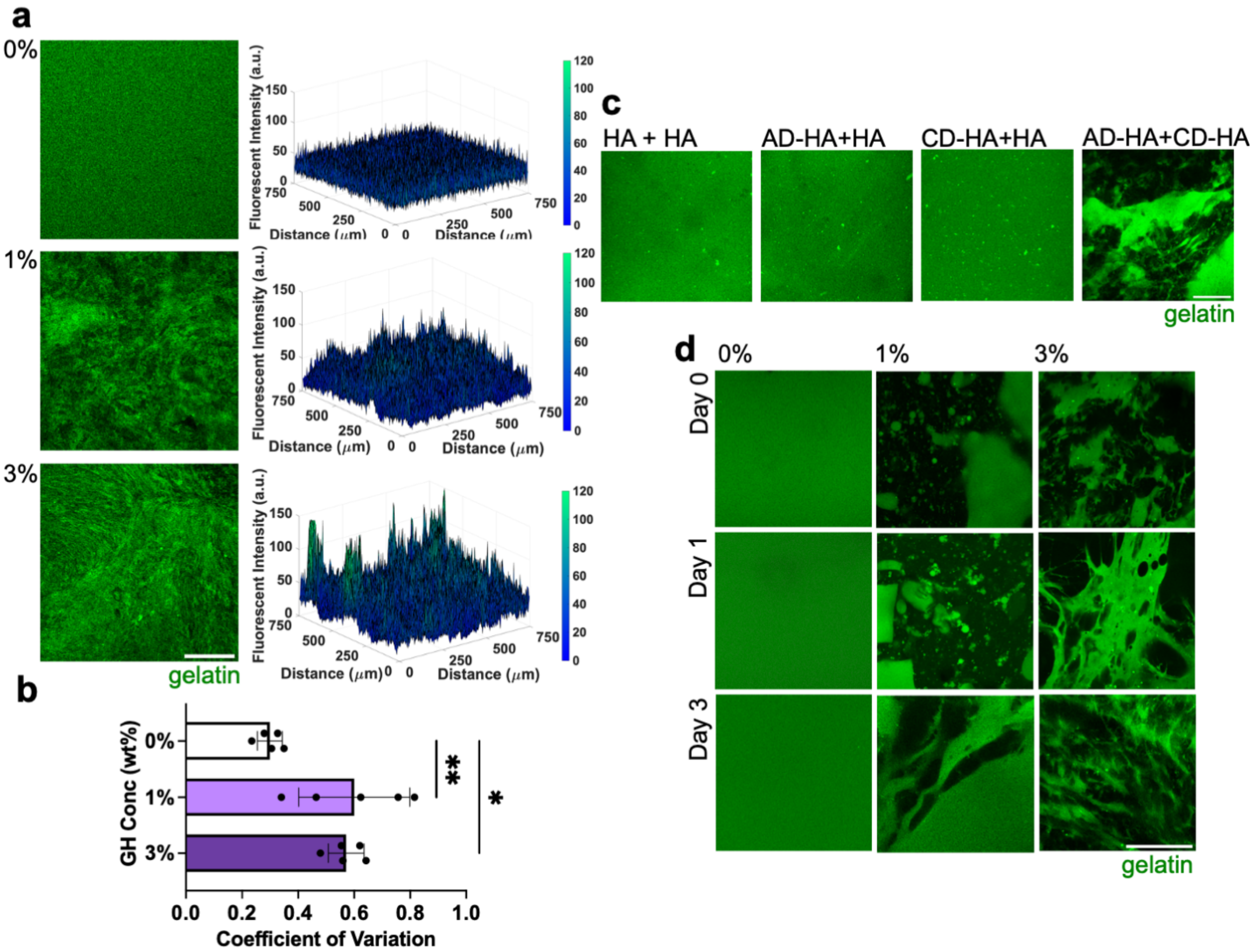


**Extended Data Fig. 2: Bicontinuous hydrogel structure relies on both GH components and is stable over time. a,** Representative single Z sections of bicontinuous hydrogels (left panel) separated into GR (green) and GP (unlabeled) domains and their corresponding fluorescent intensity profiles (right panel). Scale bar = 200 µm. **b,** Quantification of variation in structural properties of GR domains (green). n= 4-5 regions across 3 distinct gels per condition. *$p \leq .05$, **$p \leq .01$, one-way ANOVA with Tukey post hoc. **c,** Representative single Z sections of bicontinuous hydrogel structures based on presence of each GH component. Scale bar = 100 µm. **d,** Bicontinuous hydrogel structure where GR (green) and GP (unlabeled) remain distinct over 3 days. Scale bar = 100 µm. Data are mean ± s.d.

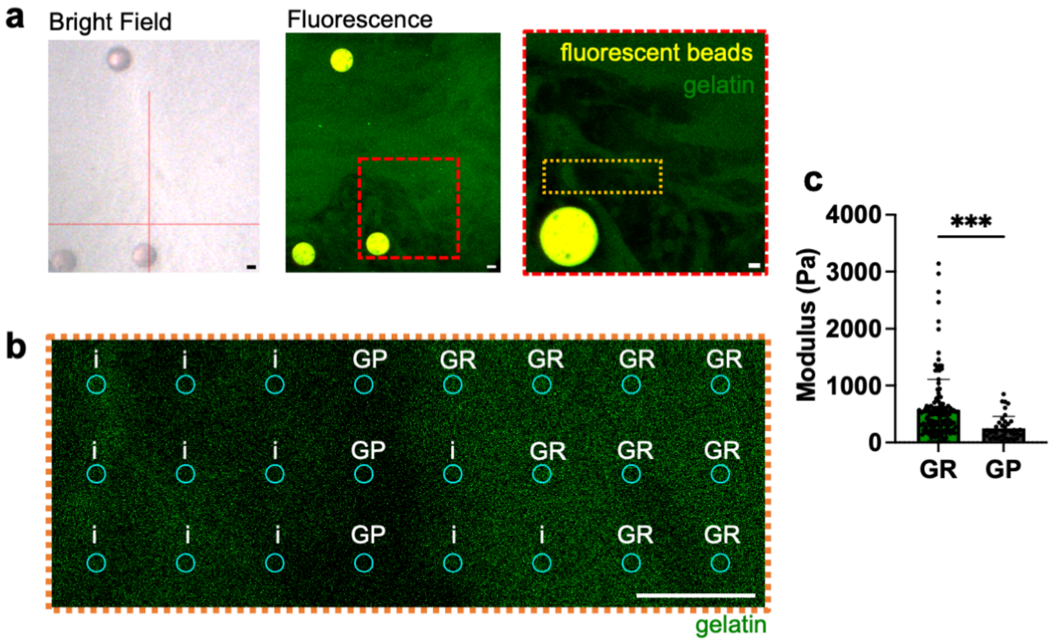


**Extended Data Fig. 3: Methodology for quantifying differential mechanical properties of GR and GP domains. a,b,** Workflow of quantification of mechanical properties in different fluorescent regions. Bright field images were taken during AFM nanoindentation (**a- left panel**, red cross represents cantilever tip; fiduciary bead-red). Fluorescent images (**a-middle panel**) of hydrogels (GR domains: green, GP domains: unlabeled) were then correlated to bright field images based on fiduciary beads (yellow). Red dashed line denotes zoom-in area corresponding to zoom-in representative fluorescent image **(a-right panel)** Orange dashed denotes further zoom-in area corresponding to analyzed grid **(b),** where indentation location is approximately denoted by cyan circles. i – inconclusive; GR – gelatin rich; GP - gelatin-poor. Scale bar = 20 $\mu$m. **c,** Elastic modulus of individual indentations based on differential fluorescent areas. n≥ 41 points across 5 hydrogels. ****p*≤.001, one-way ANOVA with Tukey post hoc. Data are mean ± s.d.

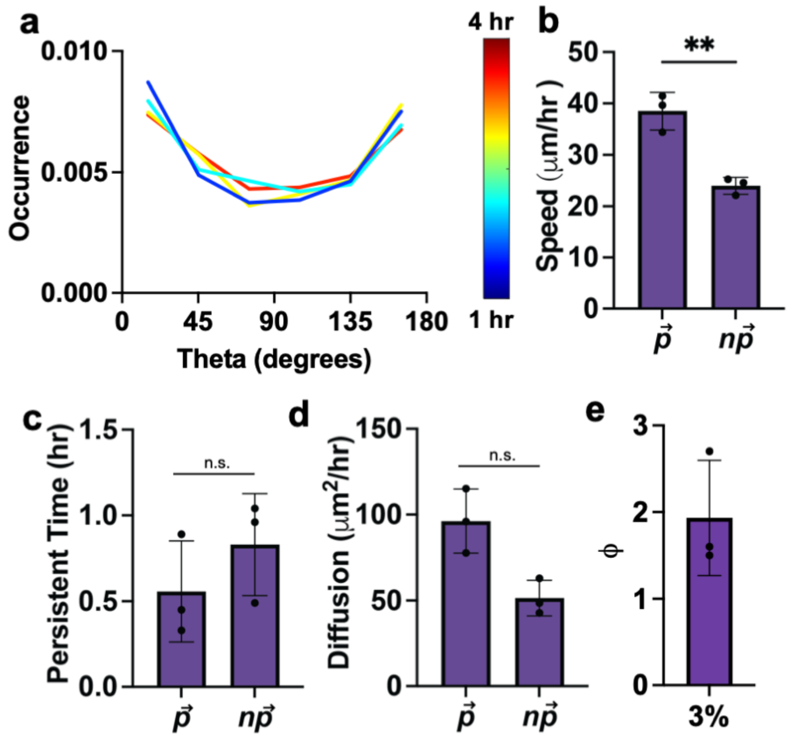


**Extended Data Fig. 4: Characterization of cell migration directionality in 3% bicontinuous hydrogel. a,** Representative distributions of angular displacements of cell outgrowth after specified time lag (denoted with heat map) of 3% GH hydrogel within one spheroid. **b-d,** Migration speed (**b**), persistence **(c),** and diffusion **(d)** along primary migration axis ($\vec{p}$) and nonprimary migration axis ($n\vec{p}$) of cells averaged per spheroid in 3% GH hydrogels. n= 3 spheroids across 1 biologically independent experiment. n.s. indicates no statistical significance, ***p*≤0.01, two-tailed paired student's t-test. **e,** Anisotropic index $\phi$ calculated from ratio of diffusion along $\vec{p}$ and $n\vec{p}$**.** n= 3 spheroids across 1 biologically independent experiment. n.s. indicates no statistical significance, two-tailed paired student's t-test. Data are mean ± s.d.

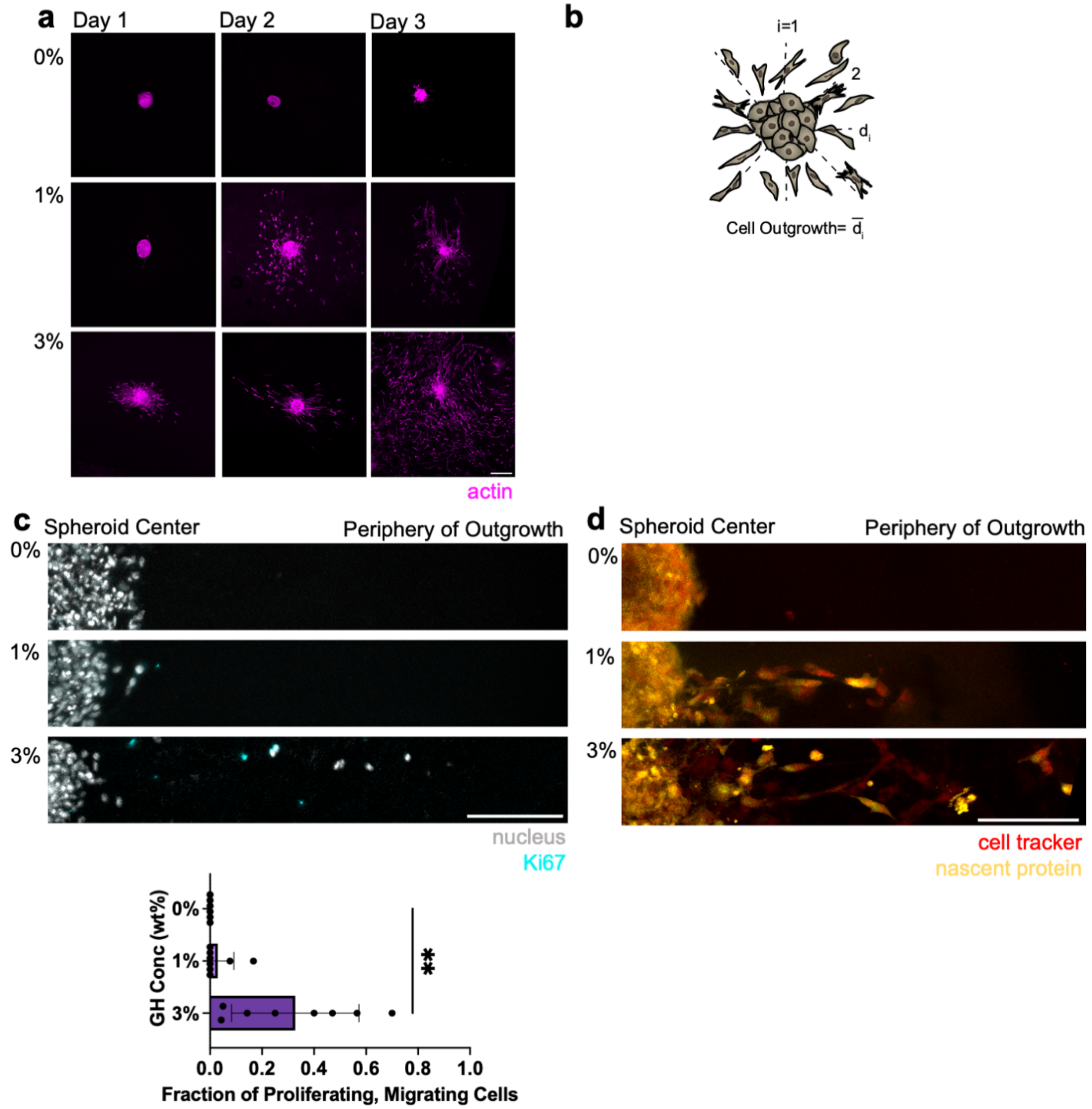


**Extended Data Fig. 5: MFC cell migration, proliferation and nascent protein deposition. a,** Representative images of cell outgrowth via actin (magenta) over time. Scale bar = 200 $\mu$m. **b,** Schematic demonstrating quantification of spheroid outgrowth. **c,** Representative images of Ki67 stain (cyan) with nuclei mask (white) over 3 days (**c-top panel,** Scale bar = 100 $\mu$m) and corresponding quantification (**c-bottom panel**). n= 6-8 spheroids per condition from 2 biologically independent experiments. **$p \leq .01$, one-way ANOVA with Tukey post hoc. **d**, Representative images of nascent protein deposition (yellow) with cell tracker (red) over 3 days. Scale bar = 100 $\mu$m. Data are mean ± s.d.

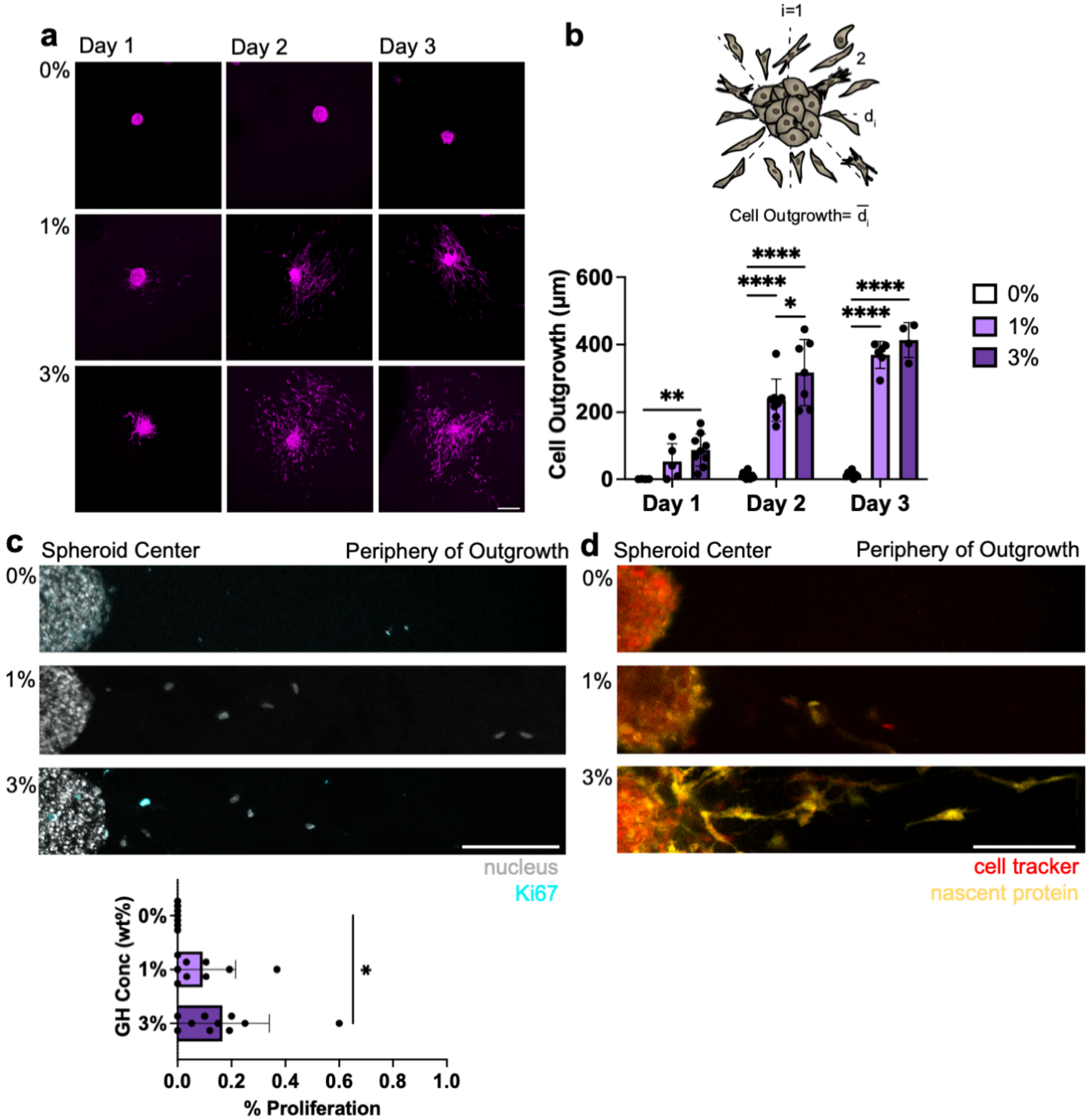


**Extended Data Fig. 6: MSC cell migration, proliferation and nascent protein deposition. a,** Representative images of cell outgrowth via actin (magenta) over 3 days. Scale bar = 200 $\mu$m. **b,** Schematic demonstrating quantification of spheroid outgrowth (top panel), and quantification of MSC cell outgrowth over time (bottom panel). n= 4-14 spheroids per condition across 2 biologically independent experiments. *$p \leq 0.05$, **$p \leq 0.01$, ****$p \leq 0.0001$, two-way ANOVA with Tukey post hoc. **c,** Representative images of Ki67 stain (cyan) with nuclei mask (white) over 3 days (**c-top panel,** Scale bar = 100 $\mu$m) and corresponding quantification (**c-bottom panel**). n= 7-10 spheroids per condition across 2 biologically independent experiments, *$p \leq .05$, one-way ANOVA with Tukey post hoc. **d**, Representative images of nascent protein deposition (yellow) with cell tracker (red) over 3 days. Scale bar = 100 $\mu$m. Data are mean ± s.d.

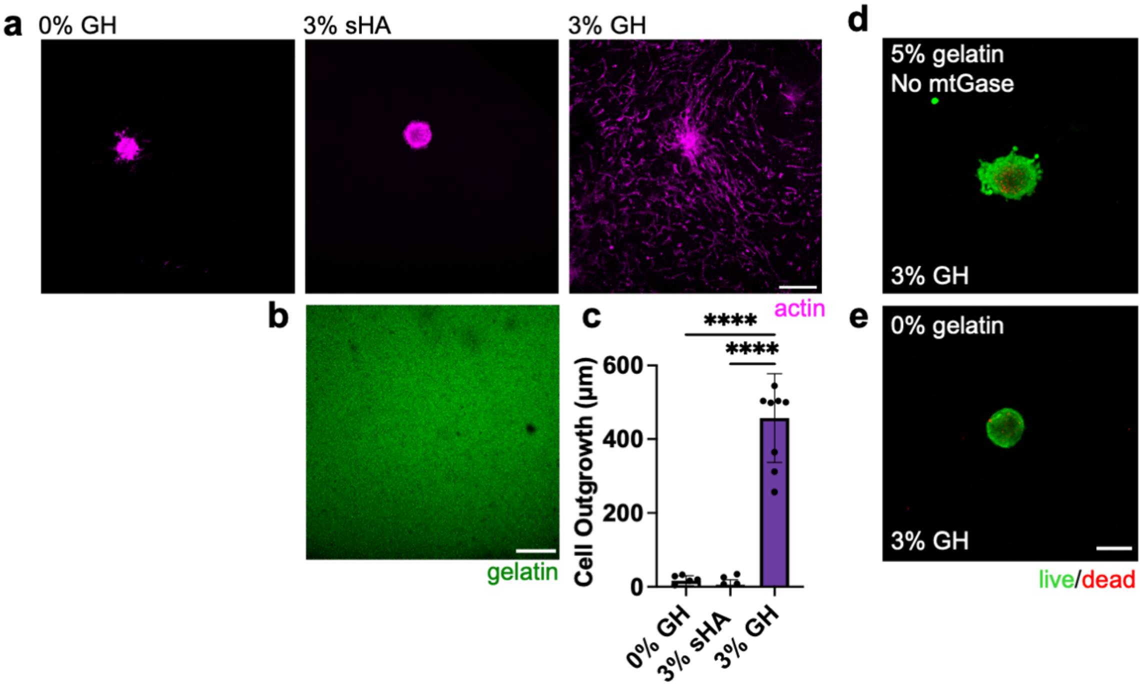


**Extended Data Fig. 7: The presence of HA alone does not account for increased migration.** **a,** Representative images of cell (actin: magenta) outgrowth at day 3 in 0 wt% GH, 3 wt% soluble HA (sHA), or 3 wt% GH (all conditions with 5 wt% gelatin and 1 U/mL transglutaminase). Scale Bar = 200 $\mu$m. **b,** Gelatin (green) distribution in hydrogels with soluble HA. Scale Bar = 200 $\mu$m. **c,** Quantification of cell outgrowth in sHA group compared to data from Fig. 3d. n= 6-10 spheroids per condition from 2 biologically independent experiments. ****$p \leq .0001$, one-way ANOVA with Tukey post hoc. **d,e,** Live (green)-Dead (red) of MFC spheroids in 3 wt% GH and 5 wt% (**d**) or 0 wt% gelatin (**e**). Scale Bar = 200 $\mu$m. Data are mean ± s.d.

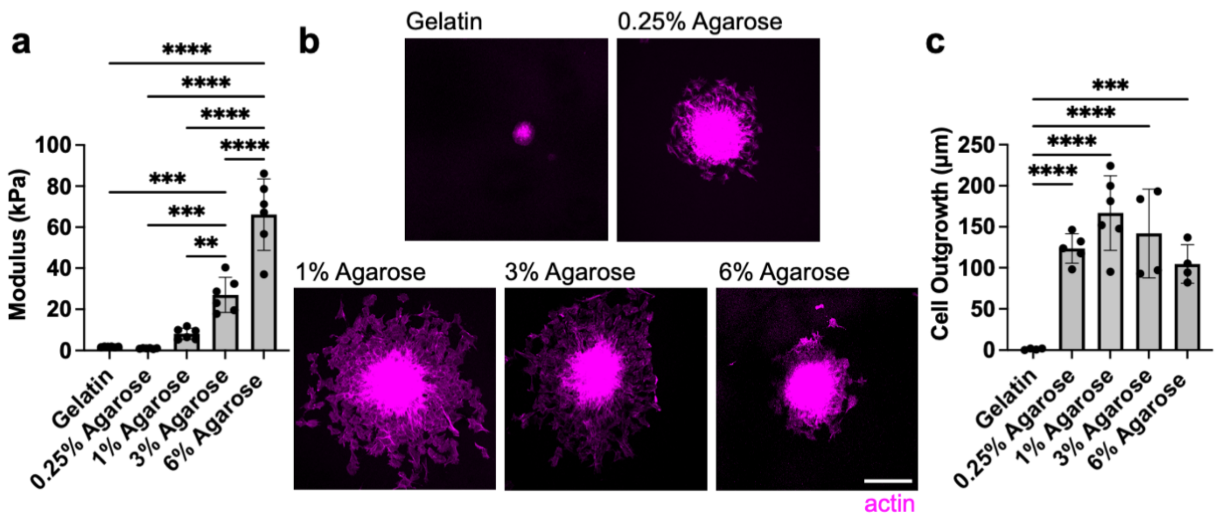


**Extended Data Fig. 8: Cells infiltrate along engineered agarose interface despite increasing differential mechanical properties across the interface. a,** Compression modulus of gelatin (5wt%, 1 U/mL enzymatic crosslinker), and varying wt% of agarose. n= 6 hydrogels per condition. $^{**}p \leq 0.01$, $^{***}p \leq 0.001$, $^{****}p \leq 0.0001$, one-way ANOVA with Tukey post hoc. **b,c,** Representative top-down images of spheroids (magenta) migrating along interface created with gelatin and top layer after 1 day (**b**, Scale bar = 200 $\mu$m) and corresponding quantification (**c**). n= 4-6 spheroids per condition across 2-3 biologically independent experiments. $^{***}p \leq 0.001$, $^{****}p \leq .0001$, one-way ANOVA with Tukey post hoc. Data are mean ± s.d.

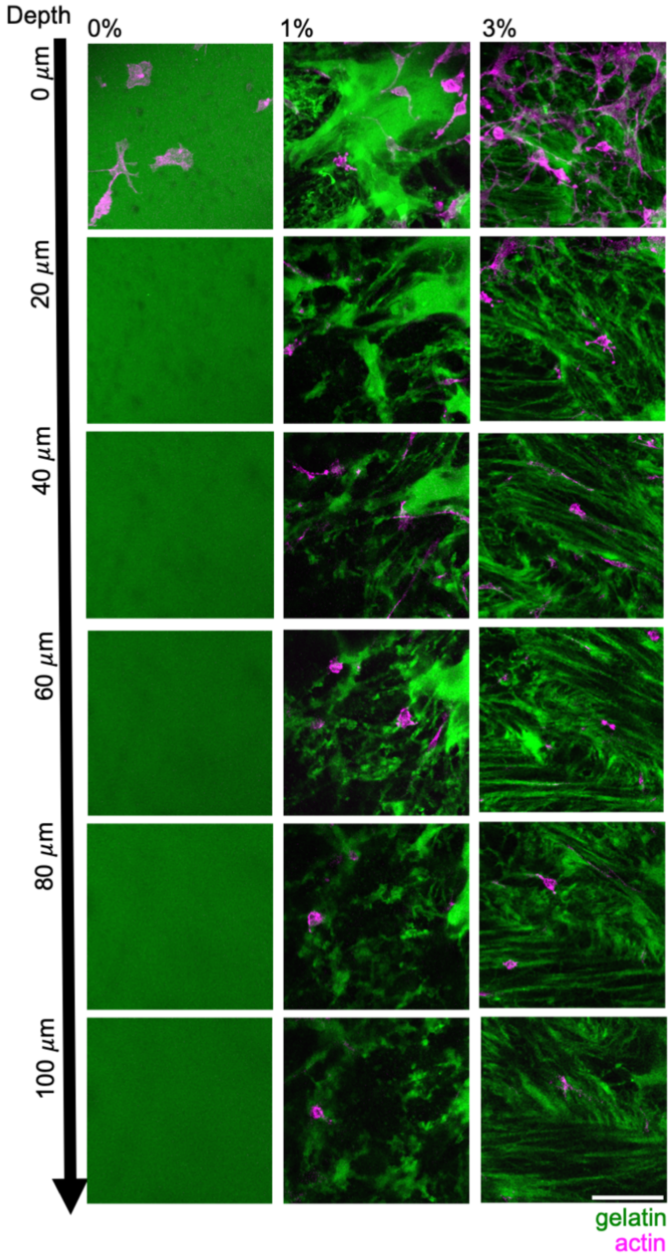


**Extended Data Fig. 9: Cell infiltration from meniscus explants are influenced by microinterfaces.** Single z-slices at different depths into the hydrogel (GR: green, GP: unlabeled) of ex vivo studies with single cell (magenta) infiltration. Scale Bar = 100 $\mu$m.

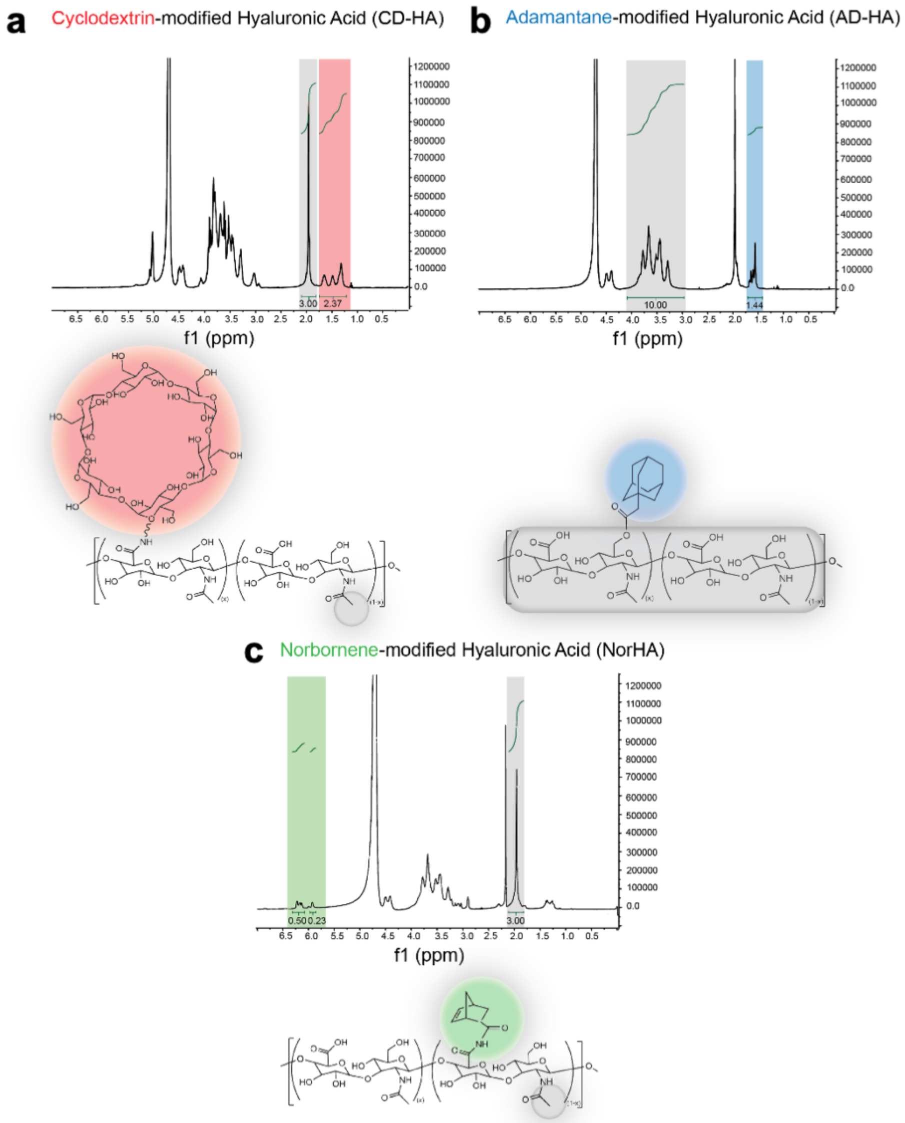


**Supplementary Fig. 1: $^{1}$H NMR spectra of synthesized CD-HA, AD-HA, and Nor-HA. a,** Chemical structure and $^{1}$H-NMR spectrum of cyclodextrin-modified hyaluronic acid (CD-HA). Modification of CD was determined by the integration of the hexane linkers (Red, 12H, δ: 1.2-1.75 ppm) when normalized to methyl group (Grey, 3H, 1.7-2.0 ppm). (**b**) Chemical structure and $^{1}$H-NMR spectrum of adamantane-modified hyaluronic acid (AD-HA). Modification of AD was determined by the integration of the ethyl multiplet (Blue, 12H, δ: 1.4-1.7 ppm) when normalized to the HA backbone (Grey, 10H, δ: 2.9-4.0 ppm). (**c**) Chemical structure and $^{1}$H-NMR spectrum of norbornene-modified hyaluronic acid (NorHA). Modification of NorHA was determined by the integration of the norbornene peaks (Green, 2H, δ: 5.8-5.9 ppm and 6.0-6.2 ppm) when normalized to the methyl group on HA (Grey, 3H, δ: 1.7-2.0 ppm).

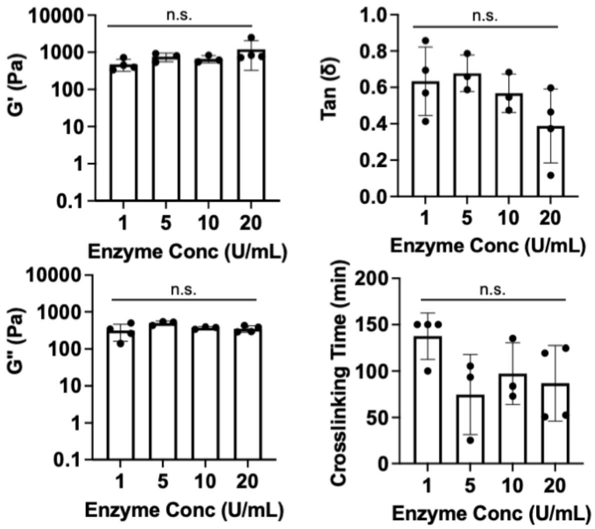


**Supplementary Fig. 2: Rheologic mechanical properties of bicontinuous hydrogels with varying enzyme concentrations.** Rheological measurements (1 Hz, 1% strain) of hydrogels (5wt% gelatin, 3wt% GH) with varying enzymatic crosslinker (1,5,10,20 U/mL) including storage modulus (top, left panel), loss modulus (bottom, left panel), tan (delta)(top, right panel), and time required for G' to reach 99% of its final modulus (bottom, right panel). n= 3-4 hydrogels per condition. n.s. indicates no statistical significance, one-way ANOVA with Tukey post hoc. Data are mean ± s.d.

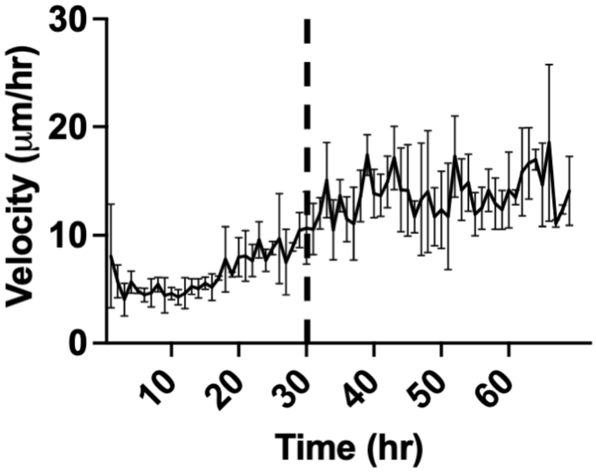


**Supplementary Fig. 3: Representative Velocity Magnitudes of Cells in 3% GH Hydrogels.** Population-averaged cell speed is time invariant once cells leave spheroid (Time ~30 hours, denoted with dashed line), a requirement for implementation of the APRW model.

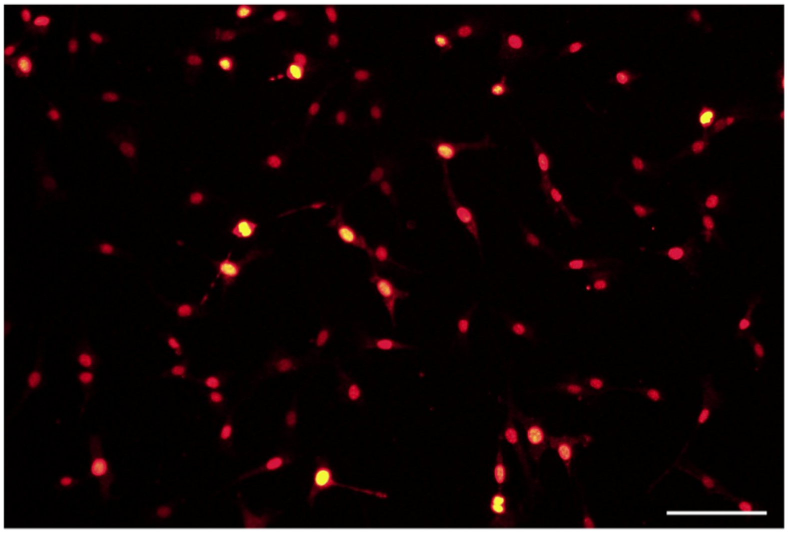

**Supplementary Fig. 4: Validation of Ki67 Antibody.** Representative image of Ki67 (red). Scale Bar = 500 $\mu$m.

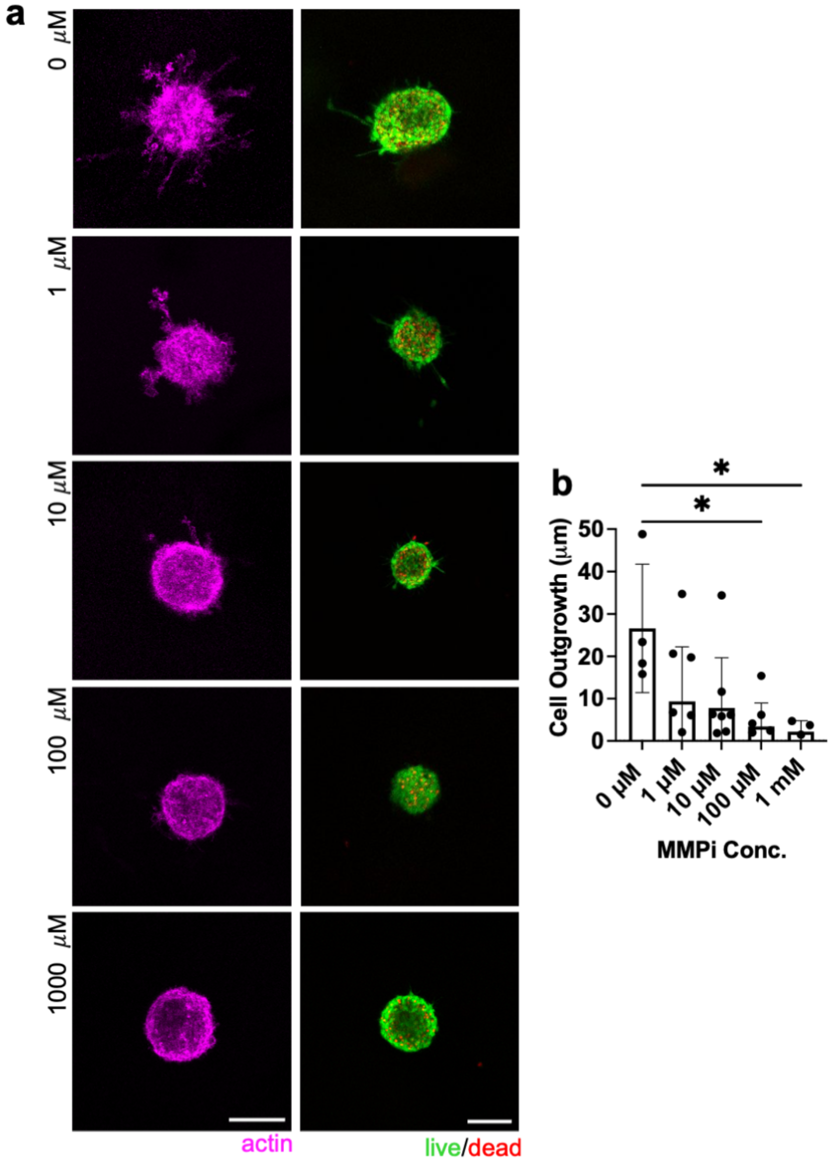


**Supplementary Fig. 5: Cell outgrowth via protease-dependent mechanisms can be tuned through MMP inhibitor concentration.** **a,** Representative images of actin (magenta) in 0% GH hydrogel after 3 days with varying Marimastat concentrations (left panels) and Live (green)-Dead (red) of MFC spheroids in corresponding inhibitor groups (right panels). Scale bar = 100 $\mu$m **b,** Quantification of cell outgrowth into 0 wt% GH hydrogels. n= 6-9 spheroids per condition from 1 biologically independent experiment. *$p \leq .05$, one-way ANOVA with Tukey post hoc. Data are mean ± s.d.

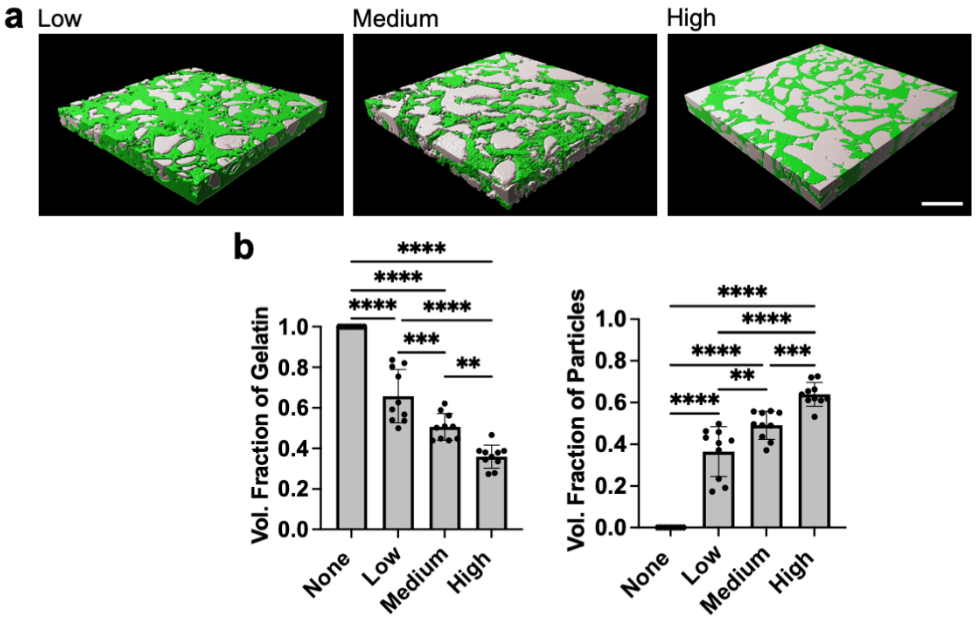


**Supplementary Fig. 6: Additional Structural Characterization of Gelatin-Agarose Particle Composite Hydrogels. a,** Representative 3D reconstructions of hydrogels with varying densification (Low, Medium, High) of agarose particles within gelatin continuous phase (gelatin: green; agarose particles: gray). Scale bar = 100 $\mu$m. b, Fraction of total volume occupied by gelatin (left panel) and agarose particles (right panel) within hydrogels. n= 9-10 regions across 3 hydrogels per condition. ***p*≤0.01, ****p*≤0.001, *****p*≤0.0001, one-way ANOVA with Tukey post hoc. Data are mean ± s.d.

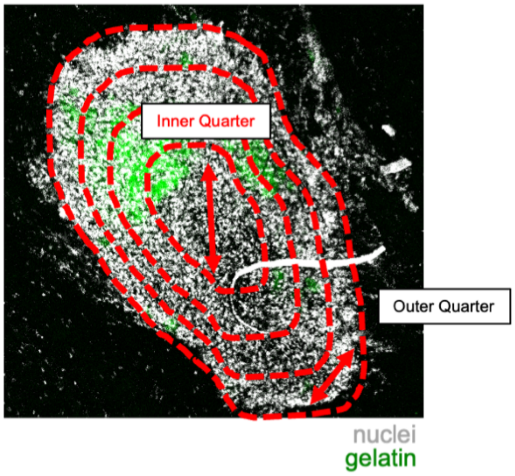


**Supplementary Fig.7: Methodology for quantifying cell density within quartiles into in vivo defect space.** Representative in vivo Z-stack of cells (nuclei: white, hydrogel: green) denoting binned areas for quantification (red dashed line).

### Supplementary Table 1

| Figure | Panel | Statistical Test | Comparison | P-value | q/t value | Degrees of freedom |
|---|---|---|---|---|---|---|
| 1 | e: left panel | One-way ANOVA Tukey Multiple Comparisons Test | 0% vs. 1% | >0.9999 | 0.002675 | 11 |
| | | | 0% vs. 3% | 0.6096 | 1.372 | 11 |
| | | | 1% vs. 3% | 0.6086 | 1.375 | 11 |
| | e: middle panel | One-way ANOVA Tukey Multiple Comparisons Test | 0% vs. 1% | 0.3813 | 1.959 | 11 |
| | | | 0% vs. 3% | 0.0001 | 9.056 | 11 |
| | | | 1% vs. 3% | 0.0009 | 7.209 | 11 |
| | e: right panel | One-way ANOVA Tukey Multiple Comparisons Test | 0% vs. 1% | 0.0325 | 4.176 | 11 |
| | | | 0% vs. 3% | <0.0001 | 13.5 | 11 |
| | | | 1% vs. 3% | <0.0001 | 9.566 | 11 |
| 2 | b: left panel | One-way ANOVA Tukey Multiple Comparisons Test | 0% vs. 1% | <0.0001 | 9.793 | 21 |
| | | | 0% vs. 3% | <0.0001 | 12.72 | 21 |
| | | | 1% vs. 3% | 0.0844 | 3.194 | 21 |
| | b: right panel | One-way ANOVA Tukey Multiple Comparisons Test | 0% vs. 1% | <0.0001 | 8.754 | 21 |
| | | | 0% vs. 3% | <0.0001 | 12.34 | 21 |
| | | | 1% vs. 3% | 0.0353 | 3.801 | 21 |
| | c: top panel | One-way ANOVA Tukey Multiple Comparisons Test | 0% vs. 1% | 0.0317 | 3.871 | 21 |
| | | | 0% vs. 3% | 0.939 | 0.4785 | 21 |
| | | | 1% vs. 3% | 0.0873 | 3.169 | 21 |
| | c: bottom panel | One-way ANOVA Tukey Multiple Comparisons Test | 0% vs. 1% | <0.0001 | 71.02 | 21 |
| | | | 0% vs. 3% | <0.0001 | 66.98 | 21 |
| | | | 1% vs. 3% | 0.5662 | 1.458 | 21 |
| | d | One-way ANOVA Tukey Multiple Comparisons Test | 0% vs. 1% | <0.0001 | 8.632 | 21 |
| | | | 0% vs. 3% | <0.0001 | 20.08 | 21 |
| | | | 1% vs. 3% | <0.0001 | 11.45 | 21 |
| | f: top panel | One-way ANOVA Tukey Multiple Comparisons Test | 0% vs. 1% | 0.0041 | 5.154 | 21 |
| | | | 0% vs. 3% | 0.0014 | 5.826 | 21 |
| | | | 1% vs. 3% | 0.9455 | 0.4515 | 21 |
| | f: bottom panel | One-way ANOVA Tukey Multiple Comparisons Test | 0% vs. 1% | 0.0354 | 3.816 | 20 |
| | | | 0% vs. 3% | 0.0228 | 4.109 | 20 |
| | | | 1% vs. 3% | 0.9792 | 0.2763 | 20 |
| | g | Two-tailed Unpaired T-test | GR vs. GP | 0.0333 | 2.315 | 17 |
| 3 | d | Two-way ANOVA Tukey Multiple Comparisons Test | Day 1:0% vs. Day 1:1% | >0.9999 | 0.2058 | 64 |
| | | | Day 1:0% vs. Day 1:3% | 0.4153 | 3.118 | 64 |
| | | | Day 1:0% vs. Day 2:0% | >0.9999 | 0.08955 | 64 |
| | | | Day 1:0% vs. Day 2:1% | 0.055 | 4.489 | 64 |
| | | | Day 1:0% vs. Day 2:3% | <0.0001 | 10.86 | 64 |
| | | | Day 1:0% vs. Day 3:0% | >0.9999 | 0.4418 | 64 |

| | | | | | | |
|---|---|---|---|---|---|---|
| | | | Day 1:0% vs. Day 3:1% | 0.0102 | 5.328 | 64 |
| | | | Day 1:0% vs. Day 3:3% | <0.0001 | 17.95 | 64 |
| | | | Day 1:1% vs. Day 1:3% | 0.4675 | 3.002 | 64 |
| | | | Day 1:1% vs. Day 2:0% | >0.9999 | 0.1198 | 64 |
| | | | Day 1:1% vs. Day 2:1% | 0.0627 | 4.416 | 64 |
| | | | Day 1:1% vs. Day 2:3% | <0.0001 | 10.94 | 64 |
| | | | Day 1:1% vs. Day 3:0% | >0.9999 | 0.2629 | 64 |
| | | | Day 1:1% vs. Day 3:1% | 0.0114 | 5.276 | 64 |
| | | | Day 1:1% vs. Day 3:3% | <0.0001 | 18.3 | 64 |
| | | | Day 1:3% vs. Day 2:0% | 0.415 | 3.119 | 64 |
| | | | Day 1:3% vs. Day 2:1% | 0.9947 | 1.202 | 64 |
| | | | Day 1:3% vs. Day 2:3% | <0.0001 | 7.974 | 64 |
| | | | Day 1:3% vs. Day 3:0% | 0.7269 | 2.445 | 64 |
| | | | Day 1:3% vs. Day 3:1% | 0.8209 | 2.21 | 64 |
| | | | Day 1:3% vs. Day 3:3% | <0.0001 | 14.75 | 64 |
| | | | Day 2:0% vs. Day 2:1% | 0.0501 | 4.539 | 64 |
| | | | Day 2:0% vs. Day 2:3% | <0.0001 | 11.05 | 64 |
| | | | Day 2:0% vs. Day 3:0% | >0.9999 | 0.3701 | 64 |
| | | | Day 2:0% vs. Day 3:1% | 0.0089 | 5.392 | 64 |
| | | | Day 2:0% vs. Day 3:3% | <0.0001 | 18.42 | 64 |
| | | | Day 2:1% vs. Day 2:3% | 0.0001 | 7.235 | 64 |
| | | | Day 2:1% vs. Day 3:0% | 0.2118 | 3.661 | 64 |
| | | | Day 2:1% vs. Day 3:1% | 0.9966 | 1.127 | 64 |
| | | | Day 2:1% vs. Day 3:3% | <0.0001 | 14.35 | 64 |
| | | | Day 2:3% vs. Day 3:0% | <0.0001 | 9.746 | 64 |
| | | | Day 2:3% vs. Day 3:1% | 0.0026 | 5.928 | 64 |
| | | | Day 2:3% vs. Day 3:3% | 0.0083 | 5.425 | 64 |
| | | | Day 3:0% vs. Day 3:1% | 0.0548 | 4.491 | 64 |
| | | | Day 3:0% vs. Day 3:3% | <0.0001 | 16.1 | 64 |
| | | | Day 3:1% vs. Day 3:3% | <0.0001 | 12.47 | 64 |
| | f | One-way ANOVA Tukey Multiple Comparisons Test | 0% vs. 1% | 0.0002 | 11.86 | 7 |
| | | | 0% vs. 3% | 0.0003 | 10.81 | 7 |
| | | | 1% vs. 3% | 0.9753 | 0.3019 | 7 |
| 4 | f | Two-way ANOVA Tukey Multiple Comparisons Test | 0%:DMSO vs. 0%:MMP inhibition | 0.999 | 0.5268 | 45 |
| | | | 0%:DMSO vs. 1%:DMSO | 0.0086 | 5.148 | 45 |
| | | | 0%:DMSO vs. 1%:MMP inhibition | 0.731 | 1.97 | 45 |
| | | | 0%:DMSO vs. 3%:DMSO | <0.0001 | 11.66 | 45 |
| | | | 0%:DMSO vs. 3%:MMP inhibition | <0.0001 | 7.412 | 45 |
| | | | 0%:MMP inhibition vs. 1%:DMSO | 0.0049 | 5.422 | 45 |

| | | | | | | |
|---|---|---|---|---|---|---|
| | | | 0%:MMP inhibition vs. 1%:MMP inhibition | 0.5452 | 2.39 | 45 |
| | | | 0%:MMP inhibition vs. 3%:DMSO | <0.0001 | 11.76 | 45 |
| | | | 0%:MMP inhibition vs. 3%:MMP inhibition | <0.0001 | 7.605 | 45 |
| | | | 1%:DMSO vs. 1%:MMP inhibition | 0.1887 | 3.352 | 45 |
| | | | 1%:DMSO vs. 3%:DMSO | 0.0005 | 6.439 | 45 |
| | | | 1%:DMSO vs. 3%:MMP inhibition | 0.9362 | 1.317 | 45 |
| | | | 1%:MMP inhibition vs. 3%:DMSO | <0.0001 | 9.951 | 45 |
| | | | 1%:MMP inhibition vs. 3%:MMP inhibition | 0.0066 | 5.278 | 45 |
| | | | 3%:DMSO vs. 3%:MMP inhibition | 0.0024 | 5.761 | 45 |
| | h | Two-tailed Unpaired T-test | sRGD vs. PBS | 0.0013 | 4.28 | 11 |
| 5 | d | One-way ANOVA Tukey Multiple Comparisons Test | No Material vs. Gelatin | <0.0001 | 15.54 | 18 |
| | | | No Material vs. Guest-Host | 0.856 | 1.124 | 18 |
| | | | No Material vs. 0.25 wt% Agarose | 0.0009 | 6.701 | 18 |
| | | | Gelatin vs. Guest-Host | <0.0001 | 14.49 | 18 |
| | | | Gelatin vs. 0.25 wt% Agarose | 0.0002 | 7.798 | 18 |
| | | | Guest-Host vs. 0.25 wt% Agarose | 0.0006 | 6.899 | 18 |
| | f: left panel | One-way ANOVA Tukey Multiple Comparisons Test | None vs. Low | >0.9999 | 0.01811 | 35 |
| | | | None vs. Medium | 0.9999 | 0.1047 | 35 |
| | | | None vs. High | 0.1085 | 3.308 | 35 |
| | | | Low vs. Medium | >0.9999 | 0.08894 | 35 |
| | | | Low vs. High | 0.0977 | 3.38 | 35 |
| | | | Medium vs. High | 0.1112 | 3.291 | 35 |
| | f: middle panel | One-way ANOVA Tukey Multiple Comparisons Test | None vs. Low | <0.0001 | 10.88 | 35 |
| | | | None vs. Medium | <0.0001 | 18.49 | 35 |
| | | | None vs. High | <0.0001 | 20.46 | 35 |
| | | | Low vs. Medium | <0.0001 | 7.813 | 35 |
| | | | Low vs. High | <0.0001 | 9.836 | 35 |
| | | | Medium vs. High | 0.4896 | 2.023 | 35 |
| | f: right panel | One-way ANOVA Tukey Multiple Comparisons Test | None vs. Low | <0.0001 | 10.28 | 35 |
| | | | None vs. Medium | <0.0001 | 13.46 | 35 |
| | | | None vs. High | <0.0001 | 16.94 | 35 |
| | | | Low vs. Medium | 0.1147 | 3.269 | 35 |
| | | | Low vs. High | 0.0001 | 6.845 | 35 |
| | | | Medium vs. High | 0.0728 | 3.576 | 35 |
| | h | One-way ANOVA Tukey Multiple Comparisons Test | None vs. Low | 0.3412 | 2.403 | 31 |
| | | | None vs. Medium | 0.0042 | 5.252 | 31 |
| | | | None vs. High | <0.0001 | 11.08 | 31 |
| | | | Low vs. Medium | 0.1967 | 2.88 | 31 |

| | | | | | | |
|---|---|---|---|---|---|---|
| | | | Low vs. High | <0.0001 | 9.376 | 31 |
| | | | Medium vs. High | <0.0001 | 7.553 | 31 |
| 6 | c: left panel | One-way ANOVA Tukey Multiple Comparisons Test | 0% vs. 1% | 0.0282 | 4.294 | 11 |
| | | | 0% vs. 3% | 0.0004 | 8.023 | 11 |
| | | | 1% vs. 3% | 0.0424 | 3.956 | 11 |
| | c: right panel | Two-way ANOVA Tukey Multiple Comparisons Test | 0-50 :0% vs. 0-50 :1% | 0.0021 | 6.519 | 55 |
| | | | 0-50 :0% vs. 0-50 :3% | <0.0001 | 11.03 | 55 |
| | | | 0-50 :0% vs. 50-100:0% | <0.0001 | 20.02 | 55 |
| | | | 0-50 :0% vs. 50-100:1% | <0.0001 | 16.14 | 55 |
| | | | 0-50 :0% vs. 50-100:3% | <0.0001 | 14.26 | 55 |
| | | | 0-50 :0% vs. 100-150:0% | <0.0001 | 20.02 | 55 |
| | | | 0-50 :0% vs. 100-150:1% | <0.0001 | 19.75 | 55 |
| | | | 0-50 :0% vs. 100-150:3% | <0.0001 | 17.95 | 55 |
| | | | 0-50 :0% vs. 150-200:0% | <0.0001 | 20.02 | 55 |
| | | | 0-50 :0% vs. 150-200:1% | <0.0001 | 20.91 | 55 |
| | | | 0-50 :0% vs. 150-200:3% | <0.0001 | 20.18 | 55 |
| | | | 0-50 :0% vs. 200+:0% | <0.0001 | 20.02 | 55 |
| | | | 0-50 :0% vs. 200+:1% | <0.0001 | 21.11 | 55 |
| | | | 0-50 :0% vs. 200+:3% | <0.0001 | 21.01 | 55 |
| | | | 0-50 :1% vs. 0-50 :3% | 0.0773 | 4.781 | 55 |
| | | | 0-50 :1% vs. 50-100:0% | <0.0001 | 14.59 | 55 |
| | | | 0-50 :1% vs. 50-100:1% | <0.0001 | 10.2 | 55 |
| | | | 0-50 :1% vs. 50-100:3% | <0.0001 | 8.206 | 55 |
| | | | 0-50 :1% vs. 100-150:0% | <0.0001 | 14.59 | 55 |
| | | | 0-50 :1% vs. 100-150:1% | <0.0001 | 14.03 | 55 |
| | | | 0-50 :1% vs. 100-150:3% | <0.0001 | 12.13 | 55 |
| | | | 0-50 :1% vs. 150-200:0% | <0.0001 | 14.59 | 55 |
| | | | 0-50 :1% vs. 150-200:1% | <0.0001 | 15.27 | 55 |
| | | | 0-50 :1% vs. 150-200:3% | <0.0001 | 14.49 | 55 |
| | | | 0-50 :1% vs. 200+:0% | <0.0001 | 14.59 | 55 |
| | | | 0-50 :1% vs. 200+:1% | <0.0001 | 15.47 | 55 |
| | | | 0-50 :1% vs. 200+:3% | <0.0001 | 15.37 | 55 |
| | | | 0-50 :3% vs. 50-100:0% | <0.0001 | 10.08 | 55 |
| | | | 0-50 :3% vs. 50-100:1% | 0.0229 | 5.42 | 55 |
| | | | 0-50 :3% vs. 50-100:3% | 0.5103 | 3.425 | 55 |
| | | | 0-50 :3% vs. 100-150:0% | <0.0001 | 10.08 | 55 |
| | | | 0-50 :3% vs. 100-150:1% | <0.0001 | 9.25 | 55 |
| | | | 0-50 :3% vs. 100-150:3% | 0.0003 | 7.345 | 55 |
| | | | 0-50 :3% vs. 150-200:0% | <0.0001 | 10.08 | 55 |
| | | | 0-50 :3% vs. 150-200:1% | <0.0001 | 10.49 | 55 |

| | | | | |
|---|---|---|---|---|
| | 0-50 :3% vs. 150-200:3% | <0.0001 | 9.711 | 55 |
| | 0-50 :3% vs. 200+:0% | <0.0001 | 10.08 | 55 |
| | 0-50 :3% vs. 200+:1% | <0.0001 | 10.69 | 55 |
| | 0-50 :3% vs. 200+:3% | <0.0001 | 10.58 | 55 |
| | 50-100:0% vs. 50-100:1% | 0.055 | 4.969 | 55 |
| | 50-100:0% vs. 50-100:3% | 0.0009 | 6.849 | 55 |
| | 50-100:0% vs. 100-150:0% | >0.9999 | 2.22E-15 | 55 |
| | 50-100:0% vs. 100-150:1% | 0.9997 | 1.357 | 55 |
| | 50-100:0% vs. 100-150:3% | 0.6429 | 3.154 | 55 |
| | 50-100:0% vs. 150-200:0% | >0.9999 | 0 | 55 |
| | 50-100:0% vs. 150-200:1% | >0.9999 | 0.1921 | 55 |
| | 50-100:0% vs. 150-200:3% | >0.9999 | 0.9231 | 55 |
| | 50-100:0% vs. 200+:0% | >0.9999 | 0 | 55 |
| | 50-100:0% vs. 200+:1% | >0.9999 | 4.69E-15 | 55 |
| | 50-100:0% vs. 200+:3% | >0.9999 | 0.09919 | 55 |
| | 50-100:1% vs. 50-100:3% | 0.9829 | 1.995 | 55 |
| | 50-100:1% vs. 100-150:0% | 0.055 | 4.969 | 55 |
| | 50-100:1% vs. 100-150:1% | 0.3271 | 3.831 | 55 |
| | 50-100:1% vs. 100-150:3% | 0.9875 | 1.925 | 55 |
| | 50-100:1% vs. 150-200:0% | 0.055 | 4.969 | 55 |
| | 50-100:1% vs. 150-200:1% | 0.0458 | 5.067 | 55 |
| | 50-100:1% vs. 150-200:3% | 0.1732 | 4.291 | 55 |
| | 50-100:1% vs. 200+:0% | 0.055 | 4.969 | 55 |
| | 50-100:1% vs. 200+:1% | 0.0309 | 5.27 | 55 |
| | 50-100:1% vs. 200+:3% | 0.0379 | 5.165 | 55 |
| | 50-100:3% vs. 100-150:0% | 0.0009 | 6.849 | 55 |
| | 50-100:3% vs. 100-150:1% | 0.0098 | 5.825 | 55 |
| | 50-100:3% vs. 100-150:3% | 0.2922 | 3.92 | 55 |
| | 50-100:3% vs. 150-200:0% | 0.0009 | 6.849 | 55 |
| | 50-100:3% vs. 150-200:1% | 0.0006 | 7.061 | 55 |
| | 50-100:3% vs. 150-200:3% | 0.0035 | 6.286 | 55 |
| | 50-100:3% vs. 200+:0% | 0.0009 | 6.849 | 55 |
| | 50-100:3% vs. 200+:1% | 0.0003 | 7.265 | 55 |
| | 50-100:3% vs. 200+:3% | 0.0004 | 7.16 | 55 |
| | 100-150:0% vs. 100-150:1% | 0.9997 | 1.357 | 55 |
| | 100-150:0% vs. 100-150:3% | 0.6429 | 3.154 | 55 |
| | 100-150:0% vs. 150-200:0% | >0.9999 | 2.22E-15 | 55 |

| | | | | | |
|---|---|---|---|---|---|
| | | 100-150:0% vs. 150-200:1% | >0.9999 | 0.1921 | 55 |
| | | 100-150:0% vs. 150-200:3% | >0.9999 | 0.9231 | 55 |
| | | 100-150:0% vs. 200+:0% | >0.9999 | 2.22E-15 | 55 |
| | | 100-150:0% vs. 200+:1% | >0.9999 | 7.03E-15 | 55 |
| | | 100-150:0% vs. 200+:3% | >0.9999 | 0.09919 | 55 |
| | | 100-150:1% vs. 100-150:3% | 0.9886 | 1.905 | 55 |
| | | 100-150:1% vs. 150-200:0% | 0.9997 | 1.357 | 55 |
| | | 100-150:1% vs. 150-200:1% | 0.9999 | 1.236 | 55 |
| | | 100-150:1% vs. 150-200:3% | >0.9999 | 0.4607 | 55 |
| | | 100-150:1% vs. 200+:0% | 0.9997 | 1.357 | 55 |
| | | 100-150:1% vs. 200+:1% | 0.9993 | 1.44 | 55 |
| | | 100-150:1% vs. 200+:3% | 0.9997 | 1.335 | 55 |
| | | 100-150:3% vs. 150-200:0% | 0.6429 | 3.154 | 55 |
| | | 100-150:3% vs. 150-200:1% | 0.6489 | 3.142 | 55 |
| | | 100-150:3% vs. 150-200:3% | 0.9323 | 2.366 | 55 |
| | | 100-150:3% vs. 200+:0% | 0.6429 | 3.154 | 55 |
| | | 100-150:3% vs. 200+:1% | 0.5493 | 3.345 | 55 |
| | | 100-150:3% vs. 200+:3% | 0.601 | 3.24 | 55 |
| | | 150-200:0% vs. 150-200:1% | >0.9999 | 0.1921 | 55 |
| | | 150-200:0% vs. 150-200:3% | >0.9999 | 0.9231 | 55 |
| | | 150-200:0% vs. 200+:0% | >0.9999 | 0 | 55 |
| | | 150-200:0% vs. 200+:1% | >0.9999 | 4.69E-15 | 55 |
| | | 150-200:0% vs. 200+:3% | >0.9999 | 0.09919 | 55 |
| | | 150-200:1% vs. 150-200:3% | >0.9999 | 0.7754 | 55 |
| | | 150-200:1% vs. 200+:0% | >0.9999 | 0.1921 | 55 |
| | | 150-200:1% vs. 200+:1% | >0.9999 | 0.2037 | 55 |
| | | 150-200:1% vs. 200+:3% | >0.9999 | 0.0985 | 55 |
| | | 150-200:3% vs. 200+:0% | >0.9999 | 0.9231 | 55 |
| | | 150-200:3% vs. 200+:1% | >0.9999 | 0.9791 | 55 |
| | | 150-200:3% vs. 200+:3% | >0.9999 | 0.8739 | 55 |
| | | 200+:0% vs. 200+:1% | >0.9999 | 4.69E-15 | 55 |
| | | 200+:0% vs. 200+:3% | >0.9999 | 0.09919 | 55 |
| | | 200+:1% vs. 200+:3% | >0.9999 | 0.1052 | 55 |
| f | Two-way ANOVA Tukey Multiple Comparisons Test | Outer Quarter:0% vs. Outer Quarter:1% | 0.9734 | 1.878 | 100 |
| | | Outer Quarter:0% vs. Outer Quarter:3% | >0.9999 | 0.4979 | 100 |
| | | Outer Quarter:0% vs. Second Quarter:0% | >0.9999 | 0.6601 | 100 |
| | | Outer Quarter:0% vs. Second Quarter:1% | >0.9999 | 0.3135 | 100 |

| | | | | |
|---|---|---|---|---|
| | Outer Quarter:0% vs. Second Quarter:3% | 0.8802 | 2.352 | 100 |
| | Outer Quarter:0% vs. Third Quarter:0% | 0.5274 | 3.164 | 100 |
| | Outer Quarter:0% vs. Third Quarter:1% | >0.9999 | 0.0429 | 100 |
| | Outer Quarter:0% vs. Third Quarter:3% | 0.8428 | 2.467 | 100 |
| | Outer Quarter:0% vs. Inner Quarter :0% | 0.2399 | 3.824 | 100 |
| | Outer Quarter:0% vs. Inner Quarter :1% | 0.9754 | 1.858 | 100 |
| | Outer Quarter:0% vs. Inner Quarter :3% | 0.3824 | 3.467 | 100 |
| | Outer Quarter:1% vs. Outer Quarter:3% | 0.8845 | 2.338 | 100 |
| | Outer Quarter:1% vs. Second Quarter:0% | 0.9995 | 1.186 | 100 |
| | Outer Quarter:1% vs. Second Quarter:1% | 0.8922 | 2.311 | 100 |
| | Outer Quarter:1% vs. Second Quarter:3% | 0.117 | 4.277 | 100 |
| | Outer Quarter:1% vs. Third Quarter:0% | 0.9969 | 1.44 | 100 |
| | Outer Quarter:1% vs. Third Quarter:1% | 0.9542 | 2.025 | 100 |
| | Outer Quarter:1% vs. Third Quarter:3% | 0.0946 | 4.397 | 100 |
| | Outer Quarter:1% vs. Inner Quarter :0% | 0.9351 | 2.132 | 100 |
| | Outer Quarter:1% vs. Inner Quarter :1% | 0.2025 | 3.939 | 100 |
| | Outer Quarter:1% vs. Inner Quarter :3% | 0.0108 | 5.443 | 100 |
| | Outer Quarter:3% vs. Second Quarter:0% | 0.9996 | 1.138 | 100 |
| | Outer Quarter:3% vs. Second Quarter:1% | >0.9999 | 0.2174 | 100 |
| | Outer Quarter:3% vs. Second Quarter:3% | 0.9805 | 1.802 | 100 |
| | Outer Quarter:3% vs. Third Quarter:0% | 0.3387 | 3.567 | 100 |
| | Outer Quarter:3% vs. Third Quarter:1% | >0.9999 | 0.4792 | 100 |
| | Outer Quarter:3% vs. Third Quarter:3% | 0.9694 | 1.914 | 100 |
| | Outer Quarter:3% vs. Inner Quarter :0% | 0.1317 | 4.208 | 100 |
| | Outer Quarter:3% vs. Inner Quarter :1% | 0.999 | 1.277 | 100 |
| | Outer Quarter:3% vs. Inner Quarter :3% | 0.665 | 2.886 | 100 |
| | Second Quarter:0% vs. Second Quarter:1% | 0.9999 | 1.006 | 100 |
| | Second Quarter:0% vs. Second Quarter:3% | 0.6127 | 2.993 | 100 |
| | Second Quarter:0% vs. Third Quarter:0% | 0.8298 | 2.504 | 100 |
| | Second Quarter:0% vs. Third Quarter:1% | >0.9999 | 0.7353 | 100 |
| | Second Quarter:0% vs. Third Quarter:3% | 0.5553 | 3.108 | 100 |
| | Second Quarter:0% vs. Inner Quarter :0% | 0.5275 | 3.164 | 100 |
| | Second Quarter:0% vs. Inner Quarter :1% | 0.8121 | 2.551 | 100 |
| | Second Quarter:0% vs. Inner Quarter :3% | 0.1554 | 4.108 | 100 |

| | | | | | | |
|---|---|---|---|---|---|---|
| | | | Second Quarter:1% vs. Second Quarter:3% | 0.9301 | 2.157 | 100 |
| | | | Second Quarter:1% vs. Third Quarter:0% | 0.3119 | 3.632 | 100 |
| | | | Second Quarter:1% vs. Third Quarter:1% | >0.9999 | 0.2853 | 100 |
| | | | Second Quarter:1% vs. Third Quarter:3% | 0.9014 | 2.277 | 100 |
| | | | Second Quarter:1% vs. Inner Quarter :0% | 0.1077 | 4.324 | 100 |
| | | | Second Quarter:1% vs. Inner Quarter :1% | 0.9913 | 1.628 | 100 |
| | | | Second Quarter:1% vs. Inner Quarter :3% | 0.4497 | 3.323 | 100 |
| | | | Second Quarter:3% vs. Third Quarter:0% | 0.0113 | 5.422 | 100 |
| | | | Second Quarter:3% vs. Third Quarter:1% | 0.8594 | 2.418 | 100 |
| | | | Second Quarter:3% vs. Third Quarter:3% | >0.9999 | 0.1119 | 100 |
| | | | Second Quarter:3% vs. Inner Quarter :0% | 0.0024 | 6.062 | 100 |
| | | | Second Quarter:3% vs. Inner Quarter :1% | >0.9999 | 0.6622 | 100 |
| | | | Second Quarter:3% vs. Inner Quarter :3% | 0.9998 | 1.084 | 100 |
| | | | Third Quarter:0% vs. Third Quarter:1% | 0.4313 | 3.361 | 100 |
| | | | Third Quarter:0% vs. Third Quarter:3% | 0.0086 | 5.537 | 100 |
| | | | Third Quarter:0% vs. Inner Quarter :0% | >0.9999 | 0.66 | 100 |
| | | | Third Quarter:0% vs. Inner Quarter :1% | 0.0197 | 5.177 | 100 |
| | | | Third Quarter:0% vs. Inner Quarter :3% | 0.0007 | 6.537 | 100 |
| | | | Third Quarter:1% vs. Third Quarter:3% | 0.8167 | 2.539 | 100 |
| | | | Third Quarter:1% vs. Inner Quarter :0% | 0.1695 | 4.053 | 100 |
| | | | Third Quarter:1% vs. Inner Quarter :1% | 0.9694 | 1.914 | 100 |
| | | | Third Quarter:1% vs. Inner Quarter :3% | 0.3315 | 3.584 | 100 |
| | | | Third Quarter:3% vs. Inner Quarter :0% | 0.0018 | 6.177 | 100 |
| | | | Third Quarter:3% vs. Inner Quarter :1% | >0.9999 | 0.7826 | 100 |
| | | | Third Quarter:3% vs. Inner Quarter :3% | >0.9999 | 0.9717 | 100 |
| | | | Inner Quarter :0% vs. Inner Quarter :1% | 0.0038 | 5.869 | 100 |
| | | | Inner Quarter :0% vs. Inner Quarter :3% | 0.0001 | 7.177 | 100 |
| | | | Inner Quarter :1% vs. Inner Quarter :3% | 0.9783 | 1.828 | 100 |

### Extended Data

| Figure | Panel | Statistical Test | Comparison | P-value | q/t value | Degrees of freedom |
|---|---|---|---|---|---|---|
| 1 | b: left panel | Two-way ANOVA Tukey Multiple Comparisons Test | 0.01:0% vs. 0.01:1% | 0.9779 | 1.796 | 36 |
| | | | 0.01:0% vs. 0.01:3% | 0.8312 | 2.48 | 36 |
| | | | 0.01:0% vs. 0.1:0% | >0.9999 | 0.3337 | 36 |
| | | | 0.01:0% vs. 0.1:1% | 0.9976 | 1.363 | 36 |

| | | | | |
|---|---|---|---|---|
| | 0.01:0% vs. 0.1:3% | 0.9676 | 1.894 | 36 |
| | 0.01:0% vs. 1:0% | >0.9999 | 0.3883 | 36 |
| | 0.01:0% vs. 1:1% | >0.9999 | 0.7485 | 36 |
| | 0.01:0% vs. 1:3% | >0.9999 | 0.3966 | 36 |
| | 0.01:0% vs. 10:0% | >0.9999 | 0.3619 | 36 |
| | 0.01:0% vs. 10:1% | >0.9999 | 0.3392 | 36 |
| | 0.01:0% vs. 10:3% | 0.0753 | 4.691 | 36 |
| | 0.01:1% vs. 0.01:3% | >0.9999 | 0.9433 | 36 |
| | 0.01:1% vs. 0.1:0% | 0.9253 | 2.148 | 36 |
| | 0.01:1% vs. 0.1:1% | >0.9999 | 0.46 | 36 |
| | 0.01:1% vs. 0.1:3% | >0.9999 | 0.3309 | 36 |
| | 0.01:1% vs. 1:0% | 0.9123 | 2.206 | 36 |
| | 0.01:1% vs. 1:1% | 0.9996 | 1.111 | 36 |
| | 0.01:1% vs. 1:3% | 0.9419 | 2.065 | 36 |
| | 0.01:1% vs. 10:0% | 0.9188 | 2.178 | 36 |
| | 0.01:1% vs. 10:1% | 0.8973 | 2.265 | 36 |
| | 0.01:1% vs. 10:3% | 0.0023 | 6.556 | 36 |
| | 0.01:3% vs. 0.1:0% | 0.708 | 2.789 | 36 |
| | 0.01:3% vs. 0.1:1% | 0.9979 | 1.342 | 36 |
| | 0.01:3% vs. 0.1:3% | >0.9999 | 0.5477 | 36 |
| | 0.01:3% vs. 1:0% | 0.6856 | 2.839 | 36 |
| | 0.01:3% vs. 1:1% | 0.9662 | 1.906 | 36 |
| | 0.01:3% vs. 1:3% | 0.75 | 2.691 | 36 |
| | 0.01:3% vs. 10:0% | 0.6964 | 2.815 | 36 |
| | 0.01:3% vs. 10:1% | 0.6558 | 2.905 | 36 |
| | 0.01:3% vs. 10:3% | 0.0017 | 6.707 | 36 |
| | 0.1:0% vs. 0.1:1% | 0.9844 | 1.714 | 36 |
| | 0.1:0% vs. 0.1:3% | 0.9129 | 2.203 | 36 |
| | 0.1:0% vs. 1:0% | >0.9999 | 0.05455 | 36 |
| | 0.1:0% vs. 1:1% | 0.9997 | 1.1 | 36 |
| | 0.1:0% vs. 1:3% | >0.9999 | 0.08764 | 36 |
| | 0.1:0% vs. 10:0% | >0.9999 | 0.0282 | 36 |
| | 0.1:0% vs. 10:1% | >0.9999 | 0.01263 | 36 |
| | 0.1:0% vs. 10:3% | 0.1222 | 4.382 | 36 |
| | 0.1:1% vs. 0.1:3% | >0.9999 | 0.7293 | 36 |
| | 0.1:1% vs. 1:0% | 0.98 | 1.772 | 36 |
| | 0.1:1% vs. 1:1% | >0.9999 | 0.6514 | 36 |
| | 0.1:1% vs. 1:3% | 0.9875 | 1.667 | 36 |
| | 0.1:1% vs. 10:0% | 0.9823 | 1.744 | 36 |
| | 0.1:1% vs. 10:1% | 0.9771 | 1.805 | 36 |

| | | | | | |
|---|---|---|---|---|---|
| | | 0.1:1% vs. 10:3% | 0.0052 | 6.157 | 36 |
| | | 0.1:3% vs. 1:0% | 0.9003 | 2.254 | 36 |
| | | 0.1:3% vs. 1:1% | 0.9985 | 1.293 | 36 |
| | | 0.1:3% vs. 1:3% | 0.9265 | 2.143 | 36 |
| | | 0.1:3% vs. 10:0% | 0.9065 | 2.229 | 36 |
| | | 0.1:3% vs. 10:1% | 0.8899 | 2.293 | 36 |
| | | 0.1:3% vs. 10:3% | 0.0051 | 6.16 | 36 |
| | | 1:0% vs. 1:1% | 0.9995 | 1.158 | 36 |
| | | 1:0% vs. 1:3% | >0.9999 | 0.03714 | 36 |
| | | 1:0% vs. 10:0% | >0.9999 | 0.02635 | 36 |
| | | 1:0% vs. 10:1% | >0.9999 | 0.07013 | 36 |
| | | 1:0% vs. 10:3% | 0.1319 | 4.331 | 36 |
| | | 1:1% vs. 1:3% | 0.9997 | 1.102 | 36 |
| | | 1:1% vs. 10:0% | 0.9996 | 1.13 | 36 |
| | | 1:1% vs. 10:1% | 0.9995 | 1.154 | 36 |
| | | 1:1% vs. 10:3% | 0.0154 | 5.593 | 36 |
| | | 1:3% vs. 10:0% | >0.9999 | 0.06154 | 36 |
| | | 1:3% vs. 10:1% | >0.9999 | 0.1033 | 36 |
| | | 1:3% vs. 10:3% | 0.2061 | 4.017 | 36 |
| | | 10:0% vs. 10:1% | >0.9999 | 0.04235 | 36 |
| | | 10:0% vs. 10:3% | 0.1271 | 4.356 | 36 |
| | | 10:1% vs. 10:3% | 0.0879 | 4.594 | 36 |
| b: middle panel | Two-way ANOVA Tukey Multiple Comparisons Test | 0.01:0% vs. 0.01:1% | >0.9999 | 0.07047 | 36 |
| | | 0.01:0% vs. 0.01:3% | >0.9999 | 0.5348 | 36 |
| | | 0.01:0% vs. 0.1:0% | >0.9999 | 0.02878 | 36 |
| | | 0.01:0% vs. 0.1:1% | >0.9999 | 0.5731 | 36 |
| | | 0.01:0% vs. 0.1:3% | 0.833 | 2.475 | 36 |
| | | 0.01:0% vs. 1:0% | >0.9999 | 0.04646 | 36 |
| | | 0.01:0% vs. 1:1% | 0.9961 | 1.444 | 36 |
| | | 0.01:0% vs. 1:3% | 0.0047 | 6.207 | 36 |
| | | 0.01:0% vs. 10:0% | >0.9999 | 0.01355 | 36 |
| | | 0.01:0% vs. 10:1% | 0.9393 | 2.079 | 36 |
| | | 0.01:0% vs. 10:3% | <0.0001 | 9.444 | 36 |
| | | 0.01:1% vs. 0.01:3% | >0.9999 | 0.4946 | 36 |
| | | 0.01:1% vs. 0.1:0% | >0.9999 | 0.1008 | 36 |
| | | 0.01:1% vs. 0.1:1% | >0.9999 | 0.5332 | 36 |
| | | 0.01:1% vs. 0.1:3% | 0.8157 | 2.523 | 36 |
| | | 0.01:1% vs. 1:0% | >0.9999 | 0.1194 | 36 |
| | | 0.01:1% vs. 1:1% | 0.9958 | 1.456 | 36 |
| | | 0.01:1% vs. 1:3% | 0.003 | 6.427 | 36 |

| | | | | |
|---|---|---|---|---|
| | 0.01:1% vs. 10:0% | >0.9999 | 0.08475 | 36 |
| | 0.01:1% vs. 10:1% | 0.9291 | 2.13 | 36 |
| | 0.01:1% vs. 10:3% | <0.0001 | 9.812 | 36 |
| | 0.01:3% vs. 0.1:0% | >0.9999 | 0.5615 | 36 |
| | 0.01:3% vs. 0.1:1% | >0.9999 | 0.03287 | 36 |
| | 0.01:3% vs. 0.1:3% | 0.9762 | 1.815 | 36 |
| | 0.01:3% vs. 1:0% | >0.9999 | 0.5778 | 36 |
| | 0.01:3% vs. 1:1% | >0.9999 | 0.7667 | 36 |
| | 0.01:3% vs. 1:3% | 0.0261 | 5.306 | 36 |
| | 0.01:3% vs. 10:0% | >0.9999 | 0.5474 | 36 |
| | 0.01:3% vs. 10:1% | 0.9978 | 1.35 | 36 |
| | 0.01:3% vs. 10:3% | <0.0001 | 8.334 | 36 |
| | 0.1:0% vs. 0.1:1% | >0.9999 | 0.6035 | 36 |
| | 0.1:0% vs. 0.1:3% | 0.8236 | 2.501 | 36 |
| | 0.1:0% vs. 1:0% | >0.9999 | 0.01768 | 36 |
| | 0.1:0% vs. 1:1% | 0.9954 | 1.474 | 36 |
| | 0.1:0% vs. 1:3% | 0.0044 | 6.234 | 36 |
| | 0.1:0% vs. 10:0% | >0.9999 | 0.01523 | 36 |
| | 0.1:0% vs. 10:1% | 0.9334 | 2.109 | 36 |
| | 0.1:0% vs. 10:3% | <0.0001 | 9.471 | 36 |
| | 0.1:1% vs. 0.1:3% | 0.9425 | 2.062 | 36 |
| | 0.1:1% vs. 1:0% | >0.9999 | 0.6221 | 36 |
| | 0.1:1% vs. 1:1% | >0.9999 | 0.9232 | 36 |
| | 0.1:1% vs. 1:3% | 0.0075 | 5.965 | 36 |
| | 0.1:1% vs. 10:0% | >0.9999 | 0.5874 | 36 |
| | 0.1:1% vs. 10:1% | 0.991 | 1.597 | 36 |
| | 0.1:1% vs. 10:3% | <0.0001 | 9.351 | 36 |
| | 0.1:3% vs. 1:0% | 0.8177 | 2.518 | 36 |
| | 0.1:3% vs. 1:1% | 0.9988 | 1.262 | 36 |
| | 0.1:3% vs. 1:3% | 0.3899 | 3.491 | 36 |
| | 0.1:3% vs. 10:0% | 0.8286 | 2.487 | 36 |
| | 0.1:3% vs. 10:1% | >0.9999 | 0.6783 | 36 |
| | 0.1:3% vs. 10:3% | 0.0025 | 6.519 | 36 |
| | 1:0% vs. 1:1% | 0.9949 | 1.493 | 36 |
| | 1:0% vs. 1:3% | 0.0043 | 6.25 | 36 |
| | 1:0% vs. 10:0% | >0.9999 | 0.03291 | 36 |
| | 1:0% vs. 10:1% | 0.9296 | 2.128 | 36 |
| | 1:0% vs. 10:3% | <0.0001 | 9.487 | 36 |
| | 1:1% vs. 1:3% | 0.0335 | 5.166 | 36 |
| | 1:1% vs. 10:0% | 0.9958 | 1.458 | 36 |

| | | | | | |
|---|---|---|---|---|---|
| | | 1:1% vs. 10:1% | >0.9999 | 0.6741 | 36 |
| | | 1:1% vs. 10:3% | <0.0001 | 8.551 | 36 |
| | | 1:3% vs. 10:0% | 0.0046 | 6.22 | 36 |
| | | 1:3% vs. 10:1% | 0.0896 | 4.582 | 36 |
| | | 1:3% vs. 10:3% | 0.5989 | 3.028 | 36 |
| | | 10:0% vs. 10:1% | 0.9366 | 2.093 | 36 |
| | | 10:0% vs. 10:3% | <0.0001 | 9.457 | 36 |
| | | 10:1% vs. 10:3% | 0.0001 | 7.967 | 36 |
| b: right panel | Two-way ANOVA Tukey Multiple Comparisons Test | 0.01:0% vs. 0.01:1% | >0.9999 | 0.3699 | 36 |
| | | 0.01:0% vs. 0.01:3% | 0.0469 | 4.973 | 36 |
| | | 0.01:0% vs. 0.1:0% | >0.9999 | 0.02747 | 36 |
| | | 0.01:0% vs. 0.1:1% | 0.9352 | 2.101 | 36 |
| | | 0.01:0% vs. 0.1:3% | <0.0001 | 12.42 | 36 |
| | | 0.01:0% vs. 1:0% | >0.9999 | 0.03737 | 36 |
| | | 0.01:0% vs. 1:1% | 0.1418 | 4.282 | 36 |
| | | 0.01:0% vs. 1:3% | <0.0001 | 12.26 | 36 |
| | | 0.01:0% vs. 10:0% | >0.9999 | 0.06563 | 36 |
| | | 0.01:0% vs. 10:1% | 0.0719 | 4.719 | 36 |
| | | 0.01:0% vs. 10:3% | <0.0001 | 8.748 | 36 |
| | | 0.01:1% vs. 0.01:3% | 0.0567 | 4.862 | 36 |
| | | 0.01:1% vs. 0.1:0% | >0.9999 | 0.3988 | 36 |
| | | 0.01:1% vs. 0.1:1% | 0.9741 | 1.836 | 36 |
| | | 0.01:1% vs. 0.1:3% | <0.0001 | 12.65 | 36 |
| | | 0.01:1% vs. 1:0% | >0.9999 | 0.4093 | 36 |
| | | 0.01:1% vs. 1:1% | 0.1716 | 4.15 | 36 |
| | | 0.01:1% vs. 1:3% | <0.0001 | 12.48 | 36 |
| | | 0.01:1% vs. 10:0% | >0.9999 | 0.3007 | 36 |
| | | 0.01:1% vs. 10:1% | 0.0853 | 4.613 | 36 |
| | | 0.01:1% vs. 10:3% | <0.0001 | 8.809 | 36 |
| | | 0.01:3% vs. 0.1:0% | 0.0449 | 4.999 | 36 |
| | | 0.01:3% vs. 0.1:1% | 0.4859 | 3.272 | 36 |
| | | 0.01:3% vs. 0.1:3% | 0.001 | 6.969 | 36 |
| | | 0.01:3% vs. 1:0% | 0.0442 | 5.008 | 36 |
| | | 0.01:3% vs. 1:1% | 0.9987 | 1.268 | 36 |
| | | 0.01:3% vs. 1:3% | 0.0014 | 6.814 | 36 |
| | | 0.01:3% vs. 10:0% | 0.052 | 4.913 | 36 |
| | | 0.01:3% vs. 10:1% | >0.9999 | 0.8666 | 36 |
| | | 0.01:3% vs. 10:3% | 0.3738 | 3.53 | 36 |
| | | 0.1:0% vs. 0.1:1% | 0.9293 | 2.13 | 36 |
| | | 0.1:0% vs. 0.1:3% | <0.0001 | 12.45 | 36 |

| | | | | | | |
|---|---|---|---|---|---|---|
| | | | 0.1:0% vs. 1:0% | >0.9999 | 0.009908 | 36 |
| | | | 0.1:0% vs. 1:1% | 0.1358 | 4.311 | 36 |
| | | | 0.1:0% vs. 1:3% | <0.0001 | 12.28 | 36 |
| | | | 0.1:0% vs. 10:0% | >0.9999 | 0.0931 | 36 |
| | | | 0.1:0% vs. 10:1% | 0.0685 | 4.748 | 36 |
| | | | 0.1:0% vs. 10:3% | <0.0001 | 8.773 | 36 |
| | | | 0.1:1% vs. 0.1:3% | <0.0001 | 11.06 | 36 |
| | | | 0.1:1% vs. 1:0% | 0.9271 | 2.14 | 36 |
| | | | 0.1:1% vs. 1:1% | 0.8839 | 2.314 | 36 |
| | | | 0.1:1% vs. 1:3% | <0.0001 | 10.89 | 36 |
| | | | 0.1:1% vs. 10:0% | 0.9478 | 2.031 | 36 |
| | | | 0.1:1% vs. 10:1% | 0.713 | 2.777 | 36 |
| | | | 0.1:1% vs. 10:3% | 0.0006 | 7.219 | 36 |
| | | | 0.1:3% vs. 1:0% | <0.0001 | 12.46 | 36 |
| | | | 0.1:3% vs. 1:1% | <0.0001 | 9.059 | 36 |
| | | | 0.1:3% vs. 1:3% | >0.9999 | 0.1548 | 36 |
| | | | 0.1:3% vs. 10:0% | <0.0001 | 12.36 | 36 |
| | | | 0.1:3% vs. 10:1% | <0.0001 | 8.658 | 36 |
| | | | 0.1:3% vs. 10:3% | 0.4122 | 3.438 | 36 |
| | | | 1:0% vs. 1:1% | 0.1338 | 4.322 | 36 |
| | | | 1:0% vs. 1:3% | <0.0001 | 12.29 | 36 |
| | | | 1:0% vs. 10:0% | >0.9999 | 0.103 | 36 |
| | | | 1:0% vs. 10:1% | 0.0674 | 4.758 | 36 |
| | | | 1:0% vs. 10:3% | <0.0001 | 8.782 | 36 |
| | | | 1:1% vs. 1:3% | <0.0001 | 8.886 | 36 |
| | | | 1:1% vs. 10:0% | 0.1568 | 4.213 | 36 |
| | | | 1:1% vs. 10:1% | >0.9999 | 0.4633 | 36 |
| | | | 1:1% vs. 10:3% | 0.0307 | 5.215 | 36 |
| | | | 1:3% vs. 10:0% | <0.0001 | 12.2 | 36 |
| | | | 1:3% vs. 10:1% | <0.0001 | 8.485 | 36 |
| | | | 1:3% vs. 10:3% | 0.4805 | 3.284 | 36 |
| | | | 10:0% vs. 10:1% | 0.0804 | 4.65 | 36 |
| | | | 10:0% vs. 10:3% | <0.0001 | 8.687 | 36 |
| | | | 10:1% vs. 10:3% | 0.0615 | 4.814 | 36 |
| | d:right panel | One-way ANOVA Tukey Multiple Comparisons Test | 0% vs. 1% | 0.8975 | 0.6324 | 6 |
| | | | 0% vs. 3% | 0.0331 | 4.819 | 6 |
| | | | 1% vs. 3% | 0.0571 | 4.187 | 6 |
| 2 | b | One-way ANOVA Tukey Multiple Comparisons Test | 0% vs. 1% | 0.0059 | 5.467 | 12 |
| | | | 0% vs. 3% | 0.0114 | 4.944 | 12 |
| | | | 1% vs. 3% | 0.9276 | 0.5239 | 12 |

| | | | | | | |
|---|---|---|---|---|---|---|
| 3 | c | Two-tailed Paired T-test | GR vs. GP | 0.0003 | 3.734 | 178 |
| 4 | a | Two-tailed Paired T-test | $\vec{p}$ vs. $n\vec{p}$ | 0.0014 | 26.78 | 2 |
| | b | Two-tailed Paired T-test | $\vec{p}$ vs. $n\vec{p}$ | 0.1601 | 2.188 | 2 |
| | d | Two-tailed Paired T-test | $\vec{p}$ vs. $n\vec{p}$ | 0.9636 | 0.05153 | 2 |
| 5 | c: lower panel | One-way ANOVA Tukey Multiple Comparisons Test | 0% vs. 1% | 0.9282 | 0.521 | 19 |
| | | | 0% vs. 3% | 0.0023 | 5.608 | 19 |
| | | | 1% vs. 3% | 0.0027 | 5.495 | 19 |
| 6 | c: lower panel | One-way ANOVA Tukey Multiple Comparisons Test | 0% vs. 1% | 0.3489 | 2.004 | 23 |
| | | | 0% vs. 3% | 0.0418 | 3.662 | 23 |
| | | | 1% vs. 3% | 0.4521 | 1.73 | 23 |
| 7 | c | One-way ANOVA Tukey Multiple Comparisons Test | 0% GH vs. 3% sHA | 0.953 | 0.4182 | 22 |
| | | | 0% GH vs. 3% GH | <0.0001 | 16.09 | 22 |
| | | | 3% sHA vs. 3% GH | <0.0001 | 18.93 | 22 |
| 8 | a | One-way ANOVA Tukey Multiple Comparisons Test | Gelatin vs. 0.25% Agarose | 0.9999 | 0.223 | 25 |
| | | | Gelatin vs. 1% Agarose | 0.7309 | 1.748 | 25 |
| | | | Gelatin vs. 3% Agarose | 0.0004 | 7.015 | 25 |
| | | | Gelatin vs. 6% Agarose | <0.0001 | 17.94 | 25 |
| | | | 0.25% Agarose vs. 1% Agarose | 0.6371 | 1.971 | 25 |
| | | | 0.25% Agarose vs. 3% Agarose | 0.0002 | 7.238 | 25 |
| | | | 0.25% Agarose vs. 6% Agarose | <0.0001 | 18.16 | 25 |
| | | | 1% Agarose vs. 3% Agarose | 0.0081 | 5.267 | 25 |
| | | | 1% Agarose vs. 6% Agarose | <0.0001 | 16.19 | 25 |
| | | | 3% Agarose vs. 6% Agarose | <0.0001 | 10.93 | 25 |
| | c | One-way ANOVA Tukey Multiple Comparisons Test | Gelatin vs. 0.25% Agarose | <0.0001 | 8.675 | 20 |
| | | | Gelatin vs. 1% Agarose | <0.0001 | 12.28 | 20 |
| | | | Gelatin vs. 3% Agarose | <0.0001 | 9.342 | 20 |
| | | | Gelatin vs. 6% Agarose | 0.0008 | 6.894 | 20 |
| | | | 0.25% Agarose vs. 1% Agarose | 0.2401 | 3.036 | 20 |
| | | | 0.25% Agarose vs. 3% Agarose | 0.9216 | 1.158 | 20 |
| | | | 0.25% Agarose vs. 6% Agarose | 0.9126 | 1.197 | 20 |
| | | | 1% Agarose vs. 3% Agarose | 0.7719 | 1.644 | 20 |
| | | | 1% Agarose vs. 6% Agarose | 0.0611 | 4.092 | 20 |
| | | | 3% Agarose vs. 6% Agarose | 0.526 | 2.235 | 20 |

### Supplementary Figures

| Figure | Panel | Statistical Test | Comparison | P-value | q/t value | Degrees of freedom |
|---|---|---|---|---|---|---|

| | | | | | | |
|---|---|---|---|---|---|---|
| 2 | top, left panel | One-way ANOVA Tukey Multiple Comparisons Test | 1 vs. 5 | 0.9398 | 1.063 | 13 |
| | | | 1 vs. 10 | 0.9851 | 0.7179 | 13 |
| | | | 1 vs. 20 | 0.2671 | 3.005 | 13 |
| | | | 1 vs. 30 | 0.9706 | 0.8657 | 13 |
| | | | 5 vs. 10 | 0.9993 | 0.3231 | 13 |
| | | | 5 vs. 20 | 0.7432 | 1.719 | 13 |
| | | | 5 vs. 30 | 0.9997 | 0.2618 | 13 |
| | | | 10 vs. 20 | 0.6035 | 2.064 | 13 |
| | | | 10 vs. 30 | >0.9999 | 0.08358 | 13 |
| | | | 20 vs. 30 | 0.5726 | 2.139 | 13 |
| | bottom, left panel | One-way ANOVA Tukey Multiple Comparisons Test | 1 vs. 5 | 0.4072 | 2.563 | 13 |
| | | | 1 vs. 10 | 0.978 | 0.7983 | 13 |
| | | | 1 vs. 20 | 0.9949 | 0.5406 | 13 |
| | | | 1 vs. 30 | 0.9543 | 0.9809 | 13 |
| | | | 5 vs. 10 | 0.7689 | 1.651 | 13 |
| | | | 5 vs. 20 | 0.6039 | 2.063 | 13 |
| | | | 5 vs. 30 | 0.1616 | 3.472 | 13 |
| | | | 10 vs. 20 | 0.9995 | 0.2978 | 13 |
| | | | 10 vs. 30 | 0.7479 | 1.706 | 13 |
| | | | 20 vs. 30 | 0.8155 | 1.521 | 13 |
| | top, right panel | One-way ANOVA Tukey Multiple Comparisons Test | 1 vs. 5 | 0.9951 | 0.5351 | 13 |
| | | | 1 vs. 10 | 0.9767 | 0.8109 | 13 |
| | | | 1 vs. 20 | 0.2038 | 3.262 | 13 |
| | | | 1 vs. 30 | 0.0684 | 4.199 | 13 |
| | | | 5 vs. 10 | 0.8955 | 1.259 | 13 |
| | | | 5 vs. 20 | 0.147 | 3.555 | 13 |
| | | | 5 vs. 30 | 0.0519 | 4.423 | 13 |
| | | | 10 vs. 20 | 0.5442 | 2.209 | 13 |
| | | | 10 vs. 30 | 0.2479 | 3.077 | 13 |
| | | | 20 vs. 30 | 0.961 | 0.9377 | 13 |
| | bottom, right panel | One-way ANOVA Tukey Multiple Comparisons Test | 1 vs. 5 | 0.174 | 3.405 | 13 |
| | | | 1 vs. 10 | 0.5536 | 2.186 | 13 |
| | | | 1 vs. 20 | 0.2763 | 2.971 | 13 |
| | | | 1 vs. 30 | 0.0087 | 5.838 | 13 |
| | | | 5 vs. 10 | 0.9239 | 1.141 | 13 |
| | | | 5 vs. 20 | 0.9894 | 0.6547 | 13 |
| | | | 5 vs. 30 | 0.6302 | 1.999 | 13 |
| | | | 10 vs. 20 | 0.9939 | 0.5652 | 13 |
| | | | 10 vs. 30 | 0.2133 | 3.219 | 13 |
| | | | 20 vs. 30 | 0.3067 | 2.867 | 13 |

| | | | | | | |
|---|---|---|---|---|---|---|
| 5 | b | One-way ANOVA Tukey Multiple Comparisons Test | 0 µM vs. 1 µM | 0.0858 | 3.765 | 28 |
| | | | 0 µM vs. 10 µM | 0.058 | 4.025 | 28 |
| | | | 0 µM vs. 100 µM | 0.0123 | 4.962 | 28 |
| | | | 0 µM vs. 1 mM | 0.0264 | 4.515 | 28 |
| | | | 1 µM vs. 10 µM | 0.9983 | 0.4163 | 28 |
| | | | 1 µM vs. 100 µM | 0.79 | 1.597 | 28 |
| | | | 1 µM vs. 1 mM | 0.8081 | 1.547 | 28 |
| | | | 10 µM vs. 100 µM | 0.925 | 1.147 | 28 |
| | | | 10 µM vs. 1 mM | 0.9157 | 1.188 | 28 |
| | | | 100 µM vs. 1 mM | 0.9998 | 0.2514 | 28 |
| 6 | b: left panel | One-way ANOVA Tukey Multiple Comparisons Test | None vs. Low | <0.0001 | 13.2 | 35 |
| | | | None vs. Medium | <0.0001 | 19.01 | 35 |
| | | | None vs. High | <0.0001 | 24.68 | 35 |
| | | | Low vs. Medium | 0.0009 | 5.967 | 35 |
| | | | Low vs. High | <0.0001 | 11.79 | 35 |
| | | | Medium vs. High | 0.0012 | 5.826 | 35 |
| | b: right panel | One-way ANOVA Tukey Multiple Comparisons Test | None vs. Low | <0.0001 | 14.91 | 35 |
| | | | None vs. Medium | <0.0001 | 20.06 | 35 |
| | | | None vs. High | <0.0001 | 26.11 | 35 |
| | | | Low vs. Medium | 0.0035 | 5.294 | 35 |
| | | | Low vs. High | <0.0001 | 11.51 | 35 |
| | | | Medium vs. High | 0.0005 | 6.217 | 35 |

**Description of Additional Supplementary Information:**

**Title:** Supplementary Movie 1| Real-time mixing of 0, 1 and 3% GH hydrogels. **Description:** Single Z-section imaged in real-time starting at 10 minutes after initial mixing (GR domains: green; GP domains: unlabeled). The first set of images correspond to 10-15 minutes after mixing. The second set of images correspond to 60-65 minutes after mixing. Each frame corresponds to 1 minute. Scale bar = 100 μm.

**Title:** Supplementary Movie 2| Visualization of Bicontinuous structure in 3D. **Description:** 3D reconstructions of confocal fluorescence stacks (spanning 100 μm) of 0, 1 and 3% GH hydrogels (GR domains: green; GP domains: grey). In 0% group, no GP regions exist. Scale bar = 50 μm.

**Title:** Supplementary Movie 3| Real-time MFC cell outgrowth and interaction with bicontinuous hydrogels. **Description:** Single Z-section of embedded MFC spheroids (GR domains: green; GP domains: unlabeled; actin: magenta) with cells infiltrating hydrogels from 8-75 hours. Each frame corresponds to 1 hour. Scale bar = 100 μm.

**Title:** Supplementary Movie 4| Visualization of Cell Outgrowth and Migration Tracks in 3D. **Description:** 3D reconstruction of confocal fluorescence stacks of real-time cell outgrowth (left) and corresponding visualization of individual cell tracks (multicolor lines) over a 72-hour culture period. Each frame corresponds to 1 hour. Scale bar = 100 μm.

**Title:** Supplementary Movie 5| Real-time cell migration with corresponding bead displacement. **Description:** Representative 5 $\mu$m maximum Z projection of a cell migrating through a 3% GH hydrogel (left, GR domain: green, actin: magenta) and corresponding particle image velocimetry (right, colored arrows with magnitude normalized per frame) over time. Each frame corresponds to 10 minutes. Scale bar = 50 $\mu$m.

**Title:** Supplementary Movie 6 | Visualization of Gelatin-Agarose Particle Composite Hydrogels in 3D. **Description:** 3D reconstructions of confocal fluorescence stacks (1000 x 1000 x 100 μm) of low, medium, and high densification of composite hydrogels (gelatin: green; agarose particles: grey). Scale bar = 100 μm.
